# Aggregation-Dependent Epitope Sequence and Modification Fingerprints of Anti-Aβ Antibodies

**DOI:** 10.1101/2025.02.26.640323

**Authors:** Talucci Ivan, Leske Timon, Klafki Hans-Wolfgang, Hassan Mohammed Mehedi, Steiert Annik, Morgado Barbara, Bothe Sebastian, van Werven Lars, Liepold Thomas, Walter Jochen, Schindelin Hermann, Wiltfang Jens, Wirths Oliver, Jahn Olaf, Maric Hans-Michael

## Abstract

A hallmark of Alzheimer’s disease (AD), the most common form of dementia, is the progressive accumulation of amyloid-beta (Aβ) peptides across distinct brain regions. Anti-Aβ antibodies (Aβ-Abs) targeting specific Aβ variants are essential tools for AD research, diagnostics, and therapy. The monoclonal antibodies Aducanumab, Lecanemab, and Donanemab have recently been approved as the first disease-modifying treatments for early AD, highlighting the clinical importance of their exact binding profiles.

In this study, we systematically characterized the binding and modification requirements of 20 Aβ-Abs, including biosimilars of Aducanumab, Lecanemab, and Donanemab, across monomeric, oligomeric, and aggregated Aβ forms. Array-based analysis of 20,000 modified Aβ peptides defined binding epitopes at single-residue resolution and revealed the impact of sequence variation, including familial AD mutations, as well as diverse post-translational modifications (PTMs). Notably, genetic variants such as H6R impaired binding of therapeutic Aβ-Abs like Aducanumab. Donanemab showed strong preference for pyroglutamate-modified AβpE3–10, while Lecanemab and Aducanumab exhibited aggregation- and sequence-context-dependent binding requirements.

Comparison of peptide binding profiles with binding of full-length and aggregated Aβ via immunoprecipitation-mass spectrometry, capillary immunoassays, Western blotting, and immunohistochemistry on AD brain tissue revealed distinct aggregation-dependent binding behaviours. The valency- and context-dependence of Aducanumab binding, together with its preference for Ser8-phosphorylated Aβ, supports a dimerization-mediated binding mechanism. For Lecanemab, our data suggest that additional structural contributions beyond the minimal N-terminal epitope are required for binding to aggregated Aβ, which remain to be fully resolved.

Together, this work provides the most comprehensive dataset to date on aggregation-dependent sequence and modification selectivity of Aβ-Abs. By integrating mutational, PTM, and aggregation contexts in a unified experimental framework, we establish a resource that enables rational selection of antibodies for research and diagnostic applications, and offers mechanistic insights that may inform the design and optimization of future therapeutic antibodies in AD.

## Introduction

Alzheimer’s disease (AD) is a neurodegenerative disorder characterized by the accumulation of amyloid-beta (Aβ) [1,2] and tau tangles in the brain, leading to cognitive decline and dementia [3]. Aβ peptides are generated under physiological conditions from Amyloid Precursor Protein (APP) by sequential proteolytic cleavages by beta- and gamma-secretase [4]. The initial β-secretase cleavage of APP results in the release of soluble sAPPβ and the generation of a 99 amino acids (AA) long carboxy (C)-terminal APP fragment (C99) with an amino (N)-terminal Asp-residue. Aβ peptides are generated by subsequent gamma-secretase cleavage of C99 [5,6].

In human cerebrospinal fluid, a conserved pattern of soluble Aβ1-37, Aβ1-38, Aβ1-39, Aβ1- 40 and Aβ1-42 has been observed [7]. Aβ peptides isolated from amyloid plaques have been shown to include several variants differing in length, their N- and C-termini, and post-translational modifications (PTMs) [8], such as cyclization of N-terminal Glu(3) or Glu(11) into pyroglutamate (pE) [9], and racemization or isomerization of Asp(1) and Asp(7) [10–13]. Furthermore, phosphorylated forms of Aβ (pSer(8) and pSer(26)) have been reported in amyloid plaques based on studies employing phospho-selective anti-Aβ antibodies (Aβ-Abs), although these findings have not yet been conclusively confirmed by mass spectrometry [14–16]. In extracellular senile plaques in the brain’s functional tissue, the parenchyma, Aβ1-42 is more prevalent than Aβ1-40, and various N- and C-terminally truncated [17–20] as well as N-terminally elongated variants [21] have been described [5,22]. In particular, those starting with a cyclized pE (3) [12] or Phe at position 4 [21,23], are highly abundant in AD brains [24]. Genetic mutations [25] further multiply Aβ amino acid sequence variability. The study of the structural polymorphisms of the Aβ peptide resulting from PTMs or mutations and the potential impact on the formation of pathophysiological oligomeric forms and fibrillar aggregates is challenging [26–30]. The majority of PTMs occur in the flexible N-terminal region of the fibrils, encompassing residues 1–16, and were proposed to trigger or accelerate the fibrillation of Aβ [31–33]. The same region is responsible for metal ion coordination which may also influence fibril formation [34]. Similarly, several pathogenic APP mutations within Aβ are associated with altered folds, higher aggregation propensities, and toxicities [35–37].

Importantly, this complexity requires molecular tools with appropriate molecular specificity and sensitivity and poses enormous challenges in designing sensitive diagnostic readouts and effective targeted therapies. Passive immunotherapy using monoclonal Aβ-Abs has been investigated extensively as an approach for AD treatment resulting in Aβ-Abs with several different Aβ selectivity profiles [38] [39]. For example, Aducanumab, a recombinant human antibody, was developed after screening human memory B-cells for reactivity towards aggregated Aβ [40,41]. Bapineuzumab, the humanized form of the murine antibody 3D6 [42] has been reported to bind to both monomeric and aggregated Aβ.

Despite extensive structural, genetic, and pre-clinical investigations, clinical trials of Aβ antibodies have yielded ambiguous outcomes. Aβ antibody binding to cerebral amyloid angiopathy (CAA) has been hypothesized to correlate with the frequency of amyloid-related imaging abnormalities with edema (ARIA-E) observed in patients treated with Aβ-based immunotherapies [38]. Aducanumab, developed by Neurimmune AG and later licensed to Biogen. was investigated in two parallel Phase 3 clinical trials: EMERGE demonstrated statistically significantly slower decline in the Clinical Dementia Rating Sum of Boxes for a high dose of Aducanumab vs. placebo, while ENGAGE failed to meet its primary endpoints [43]. Marketing and development of Aducanumab was halted in 2024 [44]. Lecanemab was designed at Uppsala University, humanized by BioArctic, and later licensed to Eisai. It showed positive results in the Clarity AD trial, and, cross-study comparison suggested that Lecanemab [45,46] has lower ARIA-E frequency than Aducanumab (13% vs 25-35%) [38], Lecanemab is now marketed as Leqembi. Eli Lilly’s Donanemab, approved in the US as Kisunla, binds to CAA fibrils in proportion to the amount of pyroglutamate-modified Aβ they present [38] [47]. These divergent outcomes highlight the importance of thoroughly characterizing each antibody’s epitope specificity, binding affinity for Aβ variants, ability to access relevant brain compartments, and possible genetic confounders (such as APOE ε4).

Aβ-Abs are the primary means for the detection and targeting of specific Aβ variants within the outlined Aβ-sequence and -modification space. The precise molecular binding epitopes of Aβ-Abs are fundamental to Aβ research. However, the immunogens used to raise and select Aβ-Abs may not represent precise and reliable predictors of the actual sequence and modification requirements. Furthermore, epitope requirements reported by different mapping methodologies may differ and may thus render a comparison difficult. Here we addressed the range and limitations of the sequence and modification specificity by side-by-side comparison in complementary assays for a panel of 20 Aβ-Abs central to AD research and therapy.

## Results and Discussion

The Aβ-Abs investigated in this study are listed in **Table 1**. The panel encompasses biosimilars of the currently FDA-approved drugs (i) Aducanumab (Biogen), initially approved for the treatment of mild cognitive impairment [44] but discontinued in 2024 (ii) Lecanemab (Biogen, Uppsala University and Eisai) [48], which in contrast to Aducanumab, has been approved via regular approval track and (iii) Donanemab (Eli Lilly) as the most recent expansion of the AD-drug arsenal [49,50]. The set of Aβ-Abs further encompasses selected antibodies directed against specific Aβ variants such as Aβ1-x with a free N-terminal-Asp(1) (3D6, 82E1, 11H3, 80C2), N-truncated Aβ starting with pE(3) (AβpE3-x) (pE3-Aβ, 2-48 and Donanemab_bs_) and phosphorylated Aβ forms (1E4E11, 5H11C10). Furthermore, the panel covers Aβ-Abs against the N-terminal region of Aβ and a central epitope, as well as antibodies selective for N-terminally elongated and truncated Aβ variants.

**Table 1.**
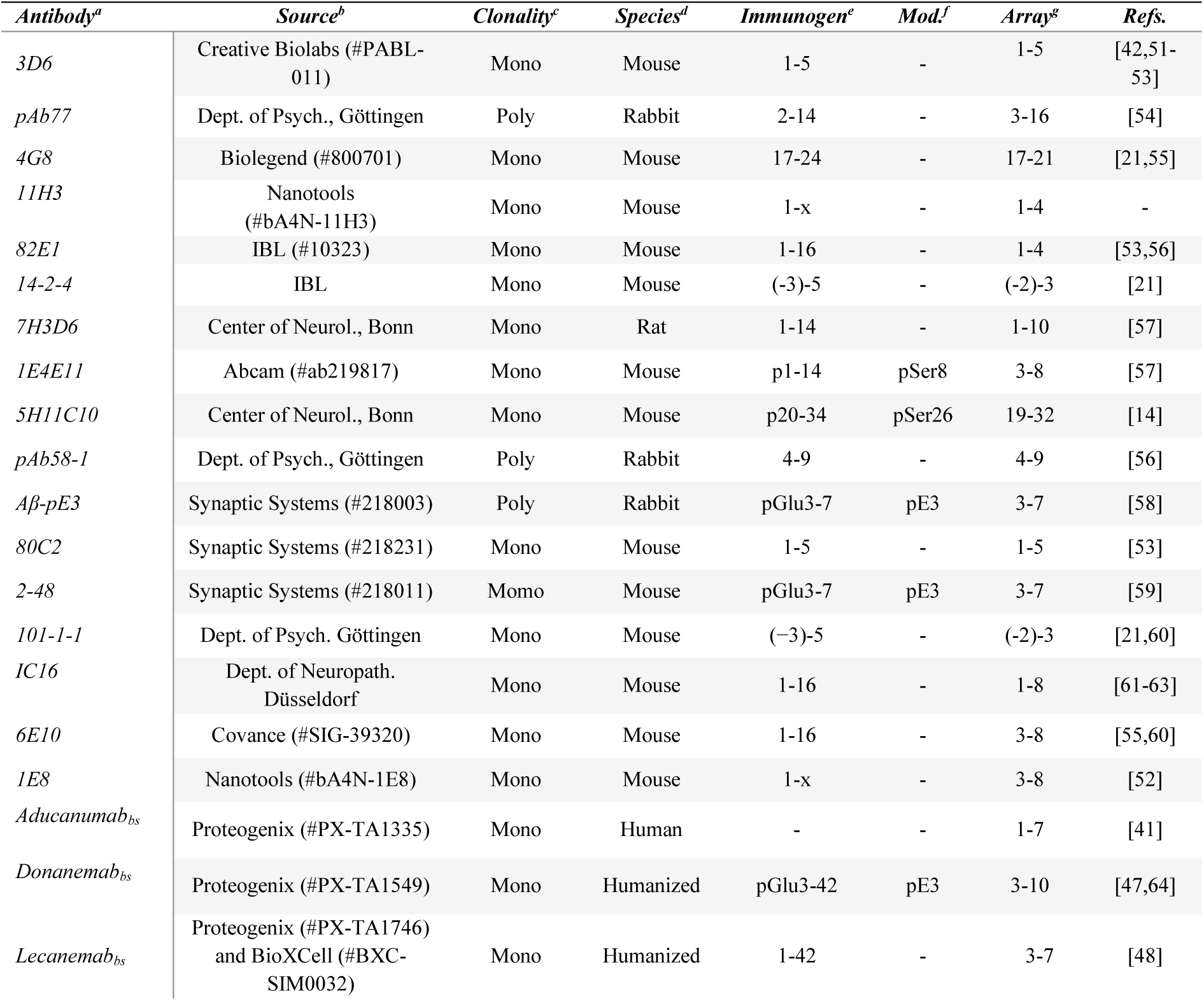
**Footnotes:** (a) names of the anti-amyloid beta antibodies. (b) Source of the antibodies used in this study, either commercial or academic. (c) Clonality of the antibodies: "mono" indicates monoclonal, while "poly" stands for polyclonal antibodies. (d) Species of origin for the IgGs: mouse, rat, human, or rabbit. (e) Immunogens, as reported in the literature, that were used to generate the corresponding antibodies. (f) If available, the anticipated PTM specificity is indicated: “p” represents phosphoserine (at positions Ser8 or Ser26), and “pE3” refers to an N-terminal pyroglutamate (at position 3). (g) epitope identified in this study on microarrays.

### Pre-screening of Aβ-Ab Epitopes

Information regarding binding epitopes and /or immunogens for most of the tested Aβ-Abs has been published and is summarized in **Table 1**. However, the binding epitope information was derived from varying methodologies. To provide an initial side-by-side comparison we pre-screened all of the listed Aβ-Abs in array format using copies of peptide libraries and standard antibody concentrations of 0.5-1 µg/mL (**Figure 1a)**. The library comprised the amyloid precursor protein (APP770) residues 649-722, corresponding to the canonical Aβ1-42 amino acid sequence with additional 23 amino terminal and 9 carboxy terminal residues. To enable single amino acid resolution, 60 peptides with a length of 15AA and overlapping by 14AA were synthesized and displayed as cellulose-conjugates incapable of oligomerization (**Figure 1b**). We first focused on Aβ-Abs with known preference for Aβ starting with a free Asp(1) and initially found that 3D6 and 82E1 also detected Aβ(2-x) and several N-terminally elongated Aβ-related peptides in addition to Aβ(1-x), at least under the generic pre-screening conditions with presumably suboptimal Ab concentrations (**Figure 1b**). Thus, it is clear that antibodies must be titrated for each specific application to obtain selectivity and to reduce non-specific binding. For optimal signal to noise ratios and ideal selectivity profiles on the microarrays, the Aβ-Ab concentrations were adjusted for 3D6, 82E1, 11H3, 80C2 and 7H3D6 (**Figure 1 – figure-supplement 1**) and (**Appendix 1-7**). This setup allowed to determine the general location of the binding epitopes within the Aβ sequence for all tested Aβ-Abs as shown by the heatmap in (**Figure 1b).** Similarly, Donanemab [47,64], 1E4E11 [57], and 5H11C10 [14], which have been previously reported as selective for AβpE3, pSer8Aβ, and pSer26-Aβ, respectively, produced detectable signals in this pre-screening with unmodified Aβ-related peptides. Again, these findings demonstrate the need for further investigation to confirm selectivity under more appropriate assay conditions, i.e. optimized antibody concentrations and presentation of modified Aβ-fragments as binding partners. The side-by-side determined epitopes are summarized together with their immunogen in **Table 1**. Epitope mapping recapitulated previously reported data, but also further confines (pAb77, 4G8, 14-2-4, 101-1- 1, 6E10, 1E8, 7H3D6, 1E4E11, 5H11C10) or even expands (Aducanumab_bs_) the required residues for binding. However, although this initial microarray only contained unmodified Aβ sequences, we also observed binding for Aβ-Abs known as PTM-selective. Specifically, Donanemab_bs_, 2-48 and Aβ-pE3 were raised against AβpE3-x (**Table 1**) and were expected to show preference for Aβ peptides starting with a pE(3). In the pre-screening with unmodified Aβ sequences, however, Donanemab_bs_ detected Aβ(3-x) starting with non-cyclized Glu (3), Aβ(-3-x) as well as several other N-terminally elongated Aβ-related peptides and Aβ1-x. Since the primary binding mode for AβpE3-x had not yet been tested at this stage these initial screening observations remain preliminary. Similarly, 1E4E11 and 5H11C10, which were reported to have high selectivity for phosphorylated Aβ peptides at Ser8 and Ser26 [14,57], both displayed strong signals with several non-phosphorylated Aβ variants in the pre- screening. Although the findings with the modification-selective Aβ-Abs are indicative of cross-reactivity due to the presence of epitopes resembling their target and/or the effect of suboptimal binding conditions, the unbiased microarray screening provided valuable initial information regarding the sequence regions and modifications of interest to be investigated in more detail in the course of our study.

**Figure 1.**
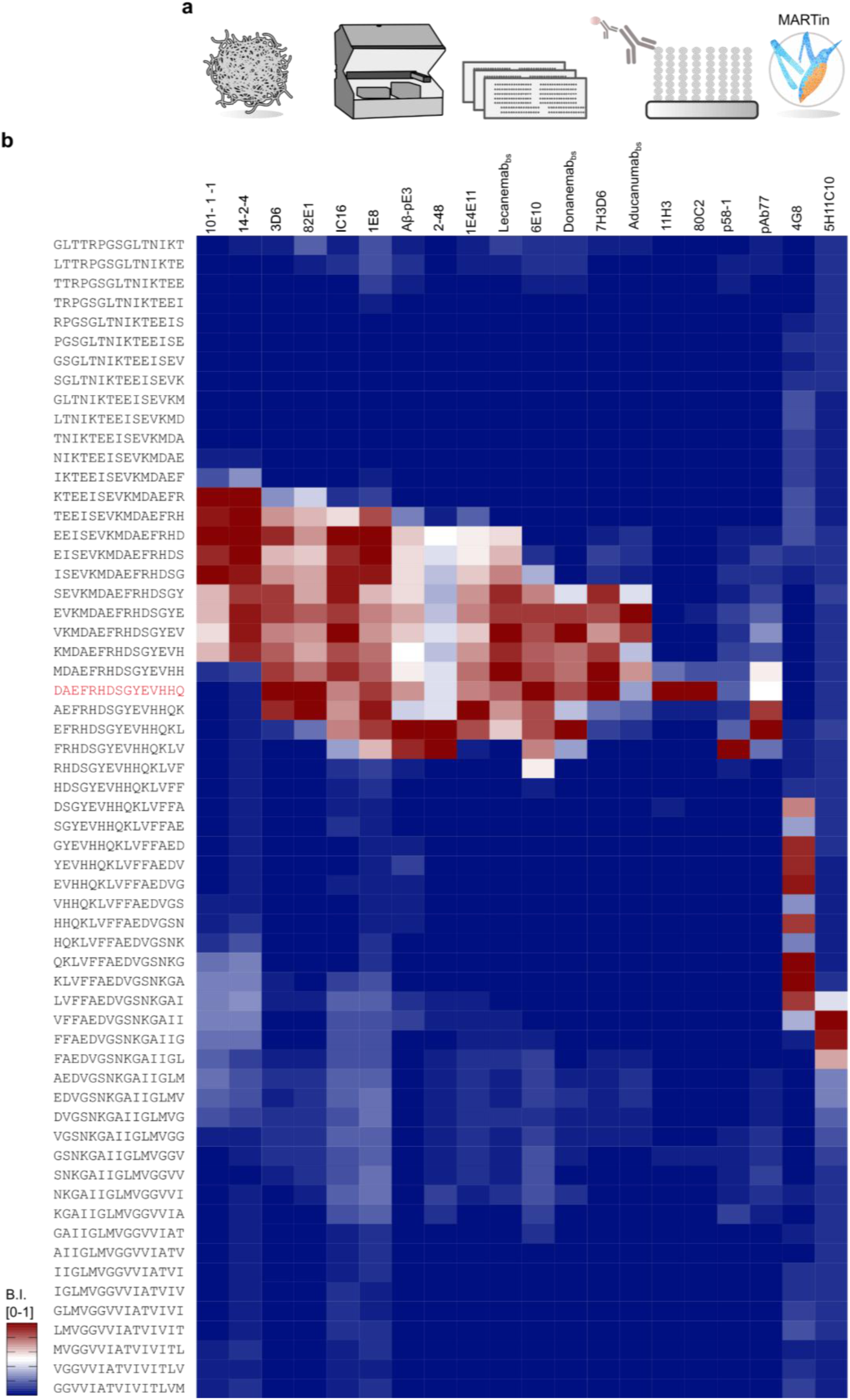
Microarray workflow and Aβ-Ab epitope pre-screening. (a) Schematic of the microarray workflow. The library comprised the amyloid precursor protein (APP) residues 649-722 (APP770 numbering), corresponding to Aβ residues -23 to 51, as a library of 15-mer peptides overlapping by 14AAs, copied in dozens, and probed for Aβ-Abs binding followed by analysis in MARTin. (b) The epitope screening binding data are summarized as a heatmap. Blue and red shades indicate the relative binding strength of the related arrayed peptides to the left and cognate Aβ-Ab on top. The blue – red scale is visualizing the normalized binding intensity for each antibody individually. The applied antibody concentrations ranged between 0.5-1 µg/mL. Note: Among the 20 tested antibodies, 1E4E11, 5H11C10, 2-48, Aβ-pE3, and Donanemab_bs_ have known PTM-selectivity, which could not be fully addressed with the initial microarray consisting of unmodified Aβ fragments only.

### Titration and Selectivity Assessment of Aβ Abs Directed Against **the Aβ N-terminus**

The sensitivity and selectivity profiles of Aβ-Abs 80C2, 11H3, 3D6, 82E1, which were expected to show preferential binding to Aβ with a free N-terminal Asp(1), and mAb 7H3D6 were explored in more detail. The canonical Aβ peptide is typically 40-42 amino acids long. In addition, various N-terminal variants have been reported [53]. Several truncated forms have been detected in brain tissue [21,24]. Notably, N-terminally truncated Aβ peptides, particularly those starting with Phe at position 4, are abundant in AD brains. The metalloproteinase ADAMTS4, which is expressed in oligodendrocytes, has been linked to APP processing and the generation of Aβ4-x peptides [23]. Additionally, N-terminally elongated Aβ peptides, such as Aβ(-3-x), which can be generated by the same enzyme in cell culture [21], may be of interest in the context of AD biomarker research: Aβ(-3-40) can serve as a reference in a composite blood-based biomarker of AD for cerebral Aβ accumulation [65,66]. Highly sensitive and selective Aβ-Abs for N-terminal Aβ variants are a prerequisite for studying their specific roles in detail. In our microarray experiments, the titration of Aβ-Abs expected to prefer Aβ1-x variants with a free N-terminal Asp (mAbs 80C2, 11H3, 3D6, 82E1) plus the rat monoclonal 7H3D6 (Figure 1 Supplement **1**) (**Appendix 1**) broadly classifies the tested mAbs into two groups. One group (80C2 and 11H3) that, on microarray under the tested experimental conditions, displayed exclusive binding towards Aβ1-x over the entire titration range albeit with limited sensitivity. A second group showed high sensitivity but concentration-dependent and limited Aβ1-x selectivity over related Aβ variants under the tested conditions (3D6, 82E1 and 7H3D6). Among the 20 tested antibodies, 1E4E11, 5H11C10, 2-48, Aβ-pE3, and Donanemab_bs_ exhibited the previously reported preferences for PTMs, which will be addressed in detail in the following paragraphs.

### Deep Mutational Scans Provide Information on Aβ-Abs Binding Requirements

Next, we further explored the reactivities observed during the epitope pre-screening (**Figure 1**) and probed in total 14 selected monoclonal antibodies with matched deep mutational libraries. Apart from the biosimilar Abs and the Abs directed against the Aβ N-terminus described above, we also selected three generic Aβ Abs as references (IC16, 1E8, 6E10) and two Abs selective for N-terminally elongated Aβ variants (14-2-4, 101-1-1), the core epitope of which we have described earlier [21]. Display and binding evaluation of all possible point-mutants systematically resolved the sequence requirements for these 14 Aβ-Abs towards the binding region corresponding to the Aβ(-10-18) amino acid sequence (**Figure 2A - figure supplement 1**) in three different 18-meric positional scans. Peptides covering Aβ(-10-8) **(Figure 2b)**, Aβ(-3-15) **(Figure 2c)** and Aβ1-18 **(Figure 2d)** regions were generated. As these three peptide regions showed binding to most of the N-terminal antibodies (**Figure 1**), we postulated that these 18-mers should cover the main antigen-antibody (Ag/Ab) contact regions. The three Aβ-related wild-type sequences were sequentially scanned from N-terminus to C-terminus by replacing each position by every proteinogenic amino acid (342 different peptide sequences in total). A customized criterion was utilized to identify the core motif for each monoclonal antibody tested. While the identified core motifs are essential for binding, they may not fully represent the specificity of the antibodies but rather indicate where the majority of the Ag/Ab contact takes place. Specifically, the raw mean intensities for all peptide variants were measured and normalized against the corresponding wildtype sequence. Amino acid exchanges that reduced binding by >50% were designated as dominant negative. This process was reiterated for each exchange at that position. Positions were considered as major contributor to the core motif when more than 50% of the exchanges were marked as dominant negative. The resulting core motifs are summarized in **Figure 2 – figure supplement 1D**. The findings highlight both similarities and differences in the binding mode of different Aβ-Abs, affirming binding to epitopes located in the N-terminal region of the Aβ sequence. Overall, this comprehensive binding data provides guidance for sorting monoclonal Aβ-Abs based on epitope properties. For two of the tested antibodies, namely 3D6 and Aducanumab, crystal structures are publicly available [41] [42], and indicate binding in the regions Aβ1-5 (3D6) and 3-7 (Aducanumab), respectively. Here, the microarray results recapitulated the epitope of 3D6 (murine precursor of Bapineuzumab) and expanded the binding interface for Aducanumab_bs_ binding to monomeric short Aβ fragments by two residues, namely Asp(1) and Ala(2) (**Figure 2D**, **Figure 2 - figure supplement 1A, B)**. Donanemab is known to recognize Aβ starting with cyclized pE3 [47,64], however, on our initial microarray screen, Donanemab_bs_ showed clear signals with AβpE3-x and several other Aβ related peptides starting N-terminally of Glu(3), including Aβ1-x (**Figure 1**). To further explore this unexpected observation, Donanemab_bs_ was tested on the Aβ (1-18) deep mutational scan. Consistent with its expected preference for the Aβ3-x region, the primary Ag/Ab contacts for the binding to Aβ1-18 involved the amino acids *EFRHDSGY.* Asp(1) and Ala(2) were not highly conserved. Importantly, the murine precursor antibody mE8 from which Donanemab was derived was previously shown to have a very strong preference for truncated AβpE3(starting with pyroglutamate) [64]. This particular point is further addressed in detail in the paragraph “Modification selectivity of monoclonal Aβ-Abs mAbs” (see below). The monoclonal Ab 1E4E11, which was previously reported to be highly selective for phosphorylated pSer(8)-Aβ [57], bound to unphosphorylated Aβ2-16 and several other unphosphorylated Aβ-related peptides on the initial screen (**Figure 1**). Interestingly, the profiling of Ser(8) did not show elevated binding to the phospho-mimetic exchanges (Asp or Glu) (**Figure 2D**). The effect of phosphorylation at Ser(8) is also addressed in detail in the paragraph “Modification selectivity of monoclonal Aβ-Abs” mAbs (see below). As shown in (**Figure 2 – figure supplement 2D**), several Aβ-Abs exhibited altered binding strength with exchanges R5G or Y10F which may be valuable information when studying the co-aggregation dynamics between exogenous (human) and endogenous (mouse) Aβ in transgenic models.

**Figure 2.**
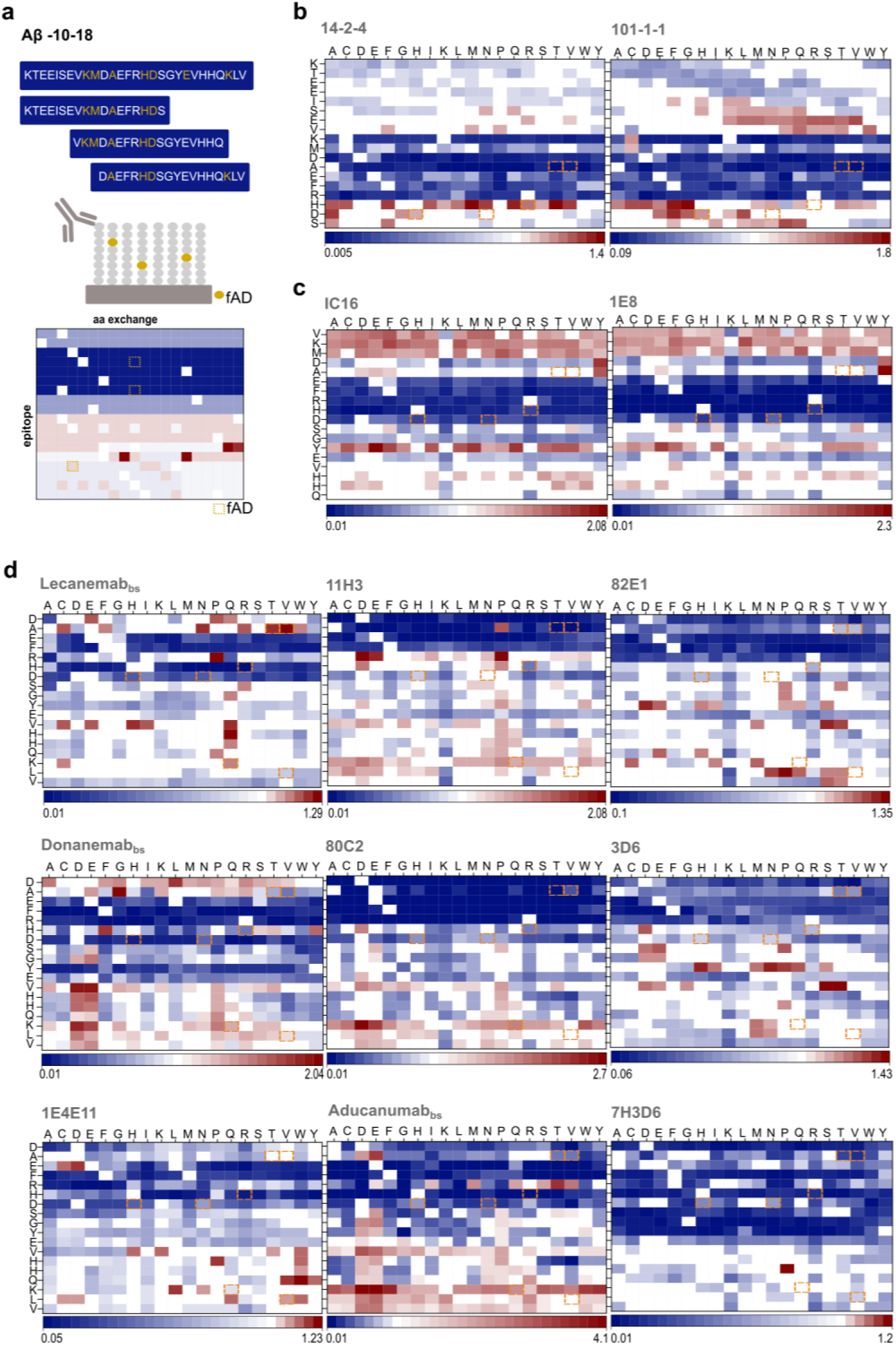
Deep Mutational Scans Define Aβ-Ab Core Sequences. (a) Schematic summary of the three deep positional scans in the N-terminal region. Seven reported APP mutations are indicated as yellow boxes. From each 18-mer epitope, positional substitutions were generated (342 variants for each epitope) and tested semi-quantitatively in microarray format. The wild-type (wt) sequence was sequentially scanned from N to C terminal by turning each position into every proteinogenic amino acid. Consequently, the contribution of each residue is depicted in blue-white-red shades, with white corresponding to no variation from wt [1] and blue-to-red representing loss or gain of binding strength. (b) Fingerprint analysis for the first peptide group covering the sequence *KTEEISEVKMDAEFRHDS*. The Aβ-Abs tested here are *101-1-1* and *14-2-4.* (c) Fingerprint analysis for the second peptide group covering the sequence *VKMDAEFRHDSGYEVHHQ*. The Aβ-Abs tested here are IC16 and 1E8. (d) Fingerprint analysis for the third peptide group covering the sequence *DAEFRHDSGYEVHHQKLV*. The Aβ-Abs tested here are Aducanumab, Donanemab and Lecanemab biosimilars. In addition, commercially available Aβ-Abs such as 3D6, 82E1, 11H3, 80C2, 7H3D6 and 1E4E11 were included. Note: The antibodies, 1E4E11 and Donanemab_bs_ have known reported PTM- preference which will be addressed in the next paragraphs.

The data set presents the sequence requirements of 14 Aβ-Abs for binding to 18-mer Aβ-related peptides in a microarray setup, thereby complementing previous structural insights and further improving our understanding of their molecular mode of action.

### Hereditary APP Mutations Affect Antibody Binding

Next, we studied the impact of some reported hereditary mutations in the *APP* gene resulting in amino acid exchanges within the Aβ sequence [25]. Specific familiar *APP* gene mutations [25,67] are reported to either lead to aggressive forms of AD or to exert a protective effect. Several mutations in the Aβ peptide have been reported to cause familial AD (fAD), with all but two having a dominant pattern of inheritance [67,68]. These alterations can lead to enhanced aggregation and increased cytotoxicity, contributing to the onset and progression of early-onset AD. The following APP mutations within the Aβ amino acid sequence were included in the microarray screen: D7H [69], D7N [70], H6R [71], A2V[72], A2T [73,74], K16Q [75,76], L17V [77].

Our analysis indicates that genetic mutations causing amino acid exchanges in or close to the Aβ sequence can affect antibody binding. Regarding the biosimilar antibodies Lecanemab_bs_, Donanemab_bs_ and Aducanumab_bs_, this may potentially have relevance for rare fAD and their therapeutic implications. The APP mutants D7H/7N, H6R and A2T/V, K16Q and L17V are highlighted in the deep mutational scan data (**Figure 2** and **Figure 2 – figure supplement 1**), as they were identified within the core motifs of the Aβ-Abs. The mutations A2V, D7H, D7N, and H6R can lead to an increased aggregation propensity of Aβ and contribute to early onset fAD [78]. Specifically, D7H (Taiwan), D7N (Tottori), and H6R (English) mutations may alter the structural features and thermodynamics of Aβ42, leading to increased β-sheet [79] content. For confirmation and for highlighting the importance of considering genetic variability for therapeutic responses, we next paradigmatically explored the effects of the D7H, D7N, H6R, A2V and A2T APP mutations [71] on Aducanumab_bs_ binding to short monomeric Aβ-related peptides on microarray.

### The Mutation H6R Abolishes Aducanumab Binding

Consistently across the two deep positional scans, Aducanumab_bs_ and 6E10 showed significantly reduced binding intensity to 18-mer peptides on the array in case of the fAD H6R mutation [71]. This aligns closely with previous findings [55], which identified *RXD* as the minimal sequence required for 6E10 binding via phage mapping (present in 53% of immunoprecipitated sequences). Our data indicate that 6E10 exhibits binding to a broader motif, *EFRHD* and shows that H6 is necessary for binding. To confirm our findings and underscore the potential significance of genetic variability in possible therapeutic strategies, we subsequently measured the binding affinity of Aducanumab_bs_ to Aβ(1-15, H6R) [71]. First, we spotted the corresponding purified peptides *DAEFR**H**DSGYEVHHQ* and *DAEFR**R**DSGYEVHHQ* (**Figure 2 – figure supplement 2a**) (**Appendix 8**) at 2-fold dilutions. The semi-quantitative on chip titration confirmed a reduced binding strength of Aducanumab_bs_ for the H6R Aβ1-15 variant. For complete quantification including kinetic evaluation, we next conducted Biolayer Interferometry (BLI) experiments (**Figure 2 – figure supplement 2b**). The BLI response of Aducanumab_bs_ for the H6R was completely abolished when compared with the wildtype sequence. These data confirm the array profiling and, in broader context, highlight a possible gap in efficacy for monoclonal Aβ-Abs in individuals with certain rare hereditary mutations associated with AD, which may have implications for personalized treatment approaches. However, whether or not the H6R amino acid replacement could impact also the binding of Aducanumab_bs_ to aggregated Aβ is still unclear.

### Modification Selectivity of Monoclonal Aβ-Abs

PTMs of Aβ can significantly impact aggregation propensity [31], clearance mechanisms, and toxicity in AD. Specific PTMs might be associated with different stages of AD progression and severity, offering potential biomarkers for diagnosis and prognosis. Targeting PTMs represents a potential therapeutic strategy to prevent Aβ aggregation and neurotoxicity, thereby slowing disease progression. *In vitro*, Aβ cyclization of N-terminal Glu into pE, and racemization/isomerization of Asp were shown to increase the aggregation propensity and neurotoxicity [12,80]. Therefore, a selected group of PTM-carrying Aβ peptides (PTM-Aβ) were incorporated into the Aβ chip to explore the potential binding preference of selected Aβ- Abs, under optimized concentrations (**Figure 3**). The antibodies were tested with both PTM- Aβ and the corresponding unmodified variants to provide a head-to-head comparison. This screening was performed in the presence of Aβ phosphorylated at positions Ser(8) or Ser(26), N-terminal pE at position 3, racemization/isomerization of Asp at position 1, and unmodified N-terminal variants. The dataset is divided into sections (**Figure 3 a-f**) for enhanced visualization, where each antibody was screened simultaneously against all peptides to locate their epitope preferences. Post-hoc mass spectrometry (MS) analysis validated the correct display of the PTM variants included in the microarray (**Appendix 1-6**). A set of antibodies without reported PTM preference was applied here as controls (**refer to Table 1**), namely 11H3, 101-1-1, IC16, 82E1 and 3D6. Antibody 6E10 has been previously shown to be unable to detect Aβ-pSer(8) and was thus also included as a control antibody [81]. On the microarray and under the tested experimental conditions, Aducanumab_bs_ and 1E4E11 (**Appendix 10**) showed preference for Aβ peptide fragments phosphorylated at Ser(8) (pSer(8)). Donanemab_bs_, 2-48, and Aβ-pE3 exhibited high selectivity for Aβ peptides starting with an N-terminal pE (**Figure 3D**) over Aβ3-x starting with a non-cyclized Glu and other Aβ variants. In contrast, mAb IC16 showed weaker binding to AβpE3-x than to Aβ3-x. Abs 7H3D6 and 11H3 did not recognize Iso-Asp(1), while 11H3 interestingly showed no affinity for D-Asp(1)-modified Aβ peptides. As expected, all of the tested antibodies, which are directed against the N-terminal Aβ region, did not show appreciable binding, and only 5H11C10 bound to the phosphorylated Ser(26) when the modification was localized centrally within Aβ19-32 (**Figure 3E, F**).

**Figure 3.**
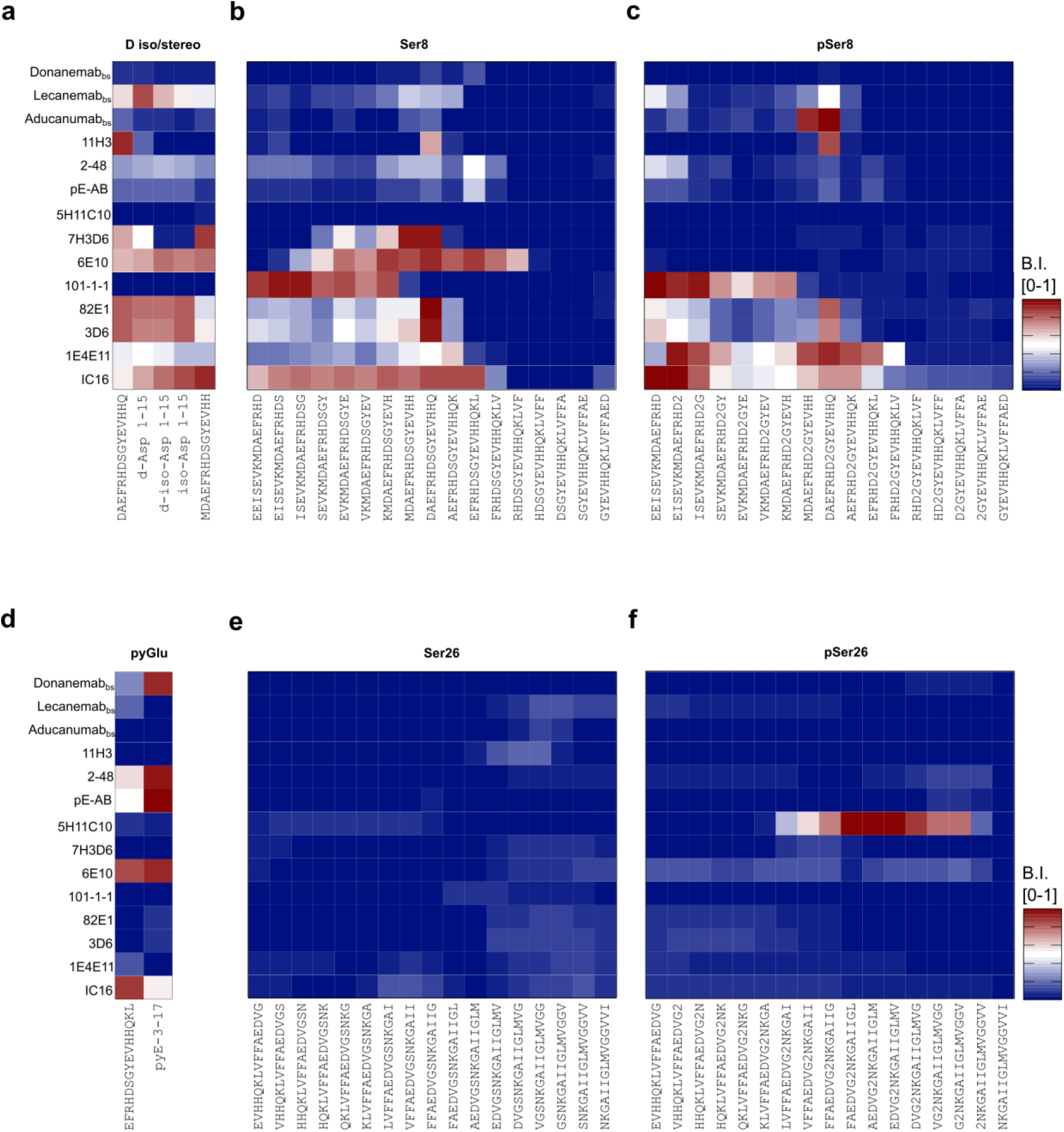
Multiplex Analysis of Aβ-Abs with preference for post-transcriptionally modified Aβ variants. The signals were normalized internally and presented as a heat map ranging from 0 to 1. Blue and red shades indicate the binding strength over the related arrayed post-transcriptionally modified peptide shown on top and cognate anti-Aβ antibody to the left. The antibodies were tested under optimal dilutions: 6E10, IC16, Aβ-pE3, 1E411, 101-1-1, 5H11C10, 2-48 at 0.25µg/mL; Lecanemab_bs_, Donanemab_bs_ at 0,0012µg/mL; Aducanumab_bs_ at 0.02µg/mL; 3D6, 11H3, 82E1 at 0.037 µg/mL; 7H3D6 at 0.012µg/mL. The dataset is split over different sections (a-f) to allow proper visualization. Each antibody was screened in parallel against all modifications. (a) The control peptide Aβ(-1-14) was used alongside Asp(1) isomers and stereoisomers. (b) The region overlapping with unmodified Serine 8. (c) The region overlapping with modified phospho Serine 8 (represented as ’2’). (d)Tthe control unmodified 3-17 was used alongside pE3. (e) The region overlapping with unmodified Serine 26. (f) The region overlapping with modified phospho Serine 26 (represented as ’2’).

### pSer8-Preference of Aducanumabbs in the Context of Non-Aggregated Aβ

Aducanumab has been reported to preferentially target aggregated Aβ [40] and bind monomeric Aβ with low micromolar affinity [41], tetrameric Aβ1-15 with low nanomolar affinity and aggregated Aβ with sub-nanomolar affinity corresponding to a roughly 10.000- fold enhanced affinity for Aβ aggregates [40,41]. However, in our hands, on microarray and under the tested experimental conditions, we observed appreciable binding of Aducanumab_bs_ to short, monomeric Aβ-related peptides (**Figures 1, 2, 3**), which we followed up by studying some molecular details. Prompted by the observed preference of Aducanumab_bs_ for phosphorylated pSer8-Aβ(-1-14) and pSer8-Aβ1-15 over unphosphorylated Aβ-related 15- mers on our microarrays (**Figure 3**,**Figure 3 - supplement 2A**), we sought to quantify the underlying relative affinity differences using the purified epitope variants in BLI as a proof-of-concept experiment with the caveat that the assay may affect the absolute affinity due to multivalency and high-density effects. As a reference, mAb 1E4E11 was titrated over the immobilized phosphorylated Aβ variant (**Figure 3, figure supplement 1**) (**Appendix 9**). As expected, mAb 1E4E11 showed a strong response in the nanomolar affinity range with fast on rates and very slow off rates, resulting in a K_D_ of 4 nM. In contrast, the overall response was at least within an order of magnitude lower with the unphosphorylated peptide and showed fast off rates, resulting in a quantified K_D_ of only 1.3μM for mAb1E4E11 binding. Overall, mAb 1E4E11 showed a 300-fold preference for the phosphorylated Aβ variant in this assay.

Henceforth, we applied this methodology to Aducanumab_bs_ (**Figure 3 – figure supplement 2**) and observed a preference for monomeric Aβ1-15 phosphorylated at Ser(8) over unphosphorylated Aβ1-15. Initially, both modified and unmodified Aβ1-15 peptides were loaded onto the BLI sensors and tested with different antibody concentrations. The affinity for the phosphorylated variant was at least 6 times higher than for the unmodified peptide, however, on microarray, the EC50 for the pSer8-Aβ1-15 was 300 times lower than that for the corresponding unphosphorylated peptide (**Appendix 10**). Importantly, this was only detectable on semi-quantitative array analysis, which can be explained by multivalency effects that may lower the observed antibody off-rate. Overall, the affinity quantification in BLI did not result in a large difference between the two variants and we thus concluded that Aducanumab_bs_ has agnostic preference for a negative charge, either provided by a carboxylic (Asp or Glu – see **Figure 2**) or phosphoserine group (**Figure 3 – figure supplement 2**), as it was not selective for the latter. This is in strong opposition to the state-of art pSer(8) antibody 1E4E11, which did not recognize phoshomimetic amino acid exchanges such as aspartate or glutamate (**Figure 2**). This observation raised the question of whether Aducanumab_bs_ is involved in electrostatic interaction.

To further explore the relevance of the observed interaction between Aducanumab_bs_ and the N-terminal phosphorylated Aβ fragment, we conducted molecular dynamics (MD) simulations (**Figure 3 – figure supplement 3**) using the published crystal structure of Aβ2-7 and Aducanumab Fab complex (PDB: 6CO3). These simulations were performed using Molecular Operating Environment (MOE) software. Specifically, we investigated two different short Aβ peptide sequences, *AEFRHDS* and *AEFRHDpS*, to examine how phosphorylation at Ser(8) affects the binding stability of the antibody-antigen complex.

By assuming that fewer fluctuations in the system correlate with a higher affinity, these data provide insights into the preferential binding mode of Aducanumab to phosphorylated short N-terminal Aβ fragments. By assuming that fewer fluctuations in the system lead to higher affinity, the root mean square deviation (RMSD) diagrams indicate greater stability (average conformation over 600ns) for the phosphorylated peptide due to its reduced fluctuations (**Figure 3 – figure supplement 3**). Additionally, the phospho group enhances the interaction with extra hydrogen bonds compared to the unmodified peptide (**Figure 3 – figure supplement 3**). Conversely, the non-phosphorylated peptide exhibits more fluctuations (**Figure 3 – figure supplement 3**). Thus, under these conditions, the MD simulation results confirm the findings from the microarray and BLI analyses.

Taken together, the MD simulations rationalize the unexpected preference of Aducanumab_bs_ towards monovalent pSer8Aβ1-15 we observed on microarray. The observation underscores the complexity of Aducanumab_bs_ binding to Aβ. This is further substantiated by comparing the previously resolved structure [41] with the MD simulation for short non-phosphorylated and pSer(8) Aβ-related peptides. Here, pSer8-Aβ2-8, unlike the corresponding non-phosphorylated Aβ, replicates the structurally determined binding mode. In the MD simulation, using the resolved epitope peptide EFRHDpS allows for the binding mode that has R317 and R318 contributing to the paratope. Similarly, the crystal structure (PDB 6CO3) includes a sulfate ion [41] that neutralizes the positive charges of R317 and R318 (**Figure 3 – figure supplement 4**). In summary, the monomeric binding mode benefits from the neutralization of a positive patch on Aducanumab Fab, which likely explains the phospho-preference.

### Immunoprecipitation with Synthetic and Endogenous Aβ-Peptides Highlights Context-Dependent Aβ-Abs Binding

Thus far, we showed that the initial microarray analysis proved invaluable in defining linear binding motifs, but needs to be complemented in different settings as more complex features of the binding behavior may be missed otherwise. In the following sections, we shift our focus to full-length Aβ peptides, which should reflect the conformations encountered *in vivo* and therefore provide insights into pathophysiologically relevant antibody binding. Thus, the screening results obtained with short Aβ fragments were next extended by analyses with intact Aβx-40 and x-42 variants. Here, the monoclonal Aβ-Abs 4G8, Donanemab_bs_, Lecanemab_bs_, Aducanumab_bs_, and 1E4E11 were tested as representative examples for their capacity in immunoprecipitating specific Aβ variants out of a mixture of ten synthetic Aβ peptides (**Figure 4**, **Appendix 11-12**). The mixture included Aβ1-40 and 1-42, N-terminal truncated Aβ(2-40, 3-40, pE3-40, 4-40, 5-40, 11-40), N-terminal elongated Aβ(-3-40) and phosphorylated pSer8- Aβ1-40 (amidated C-terminal carboxyl group). The antibodies were covalently coupled to magnetic beads and incubated, separately, with aliquots of the Aβ peptide mixture. After washing the magnetic bead immune complexes, the bound Aβ peptides were eluted, and the immunoprecipitated products were analyzed using MALDI-TOF-MS on TOPAC matrix (**Figure 4A**). As previously established [82], the TOPAC matrix allows for improved detection of phosphorylated Aβ with minimal phosphate loss. Two control antibodies, 4G8 (core epitope 17-21 according to our microarray data) and 1E4E11, were used to set up the system, representing respectively, “pan-Abeta” and pSer8-Aβ1-40 profiles.

**Figure 4:**
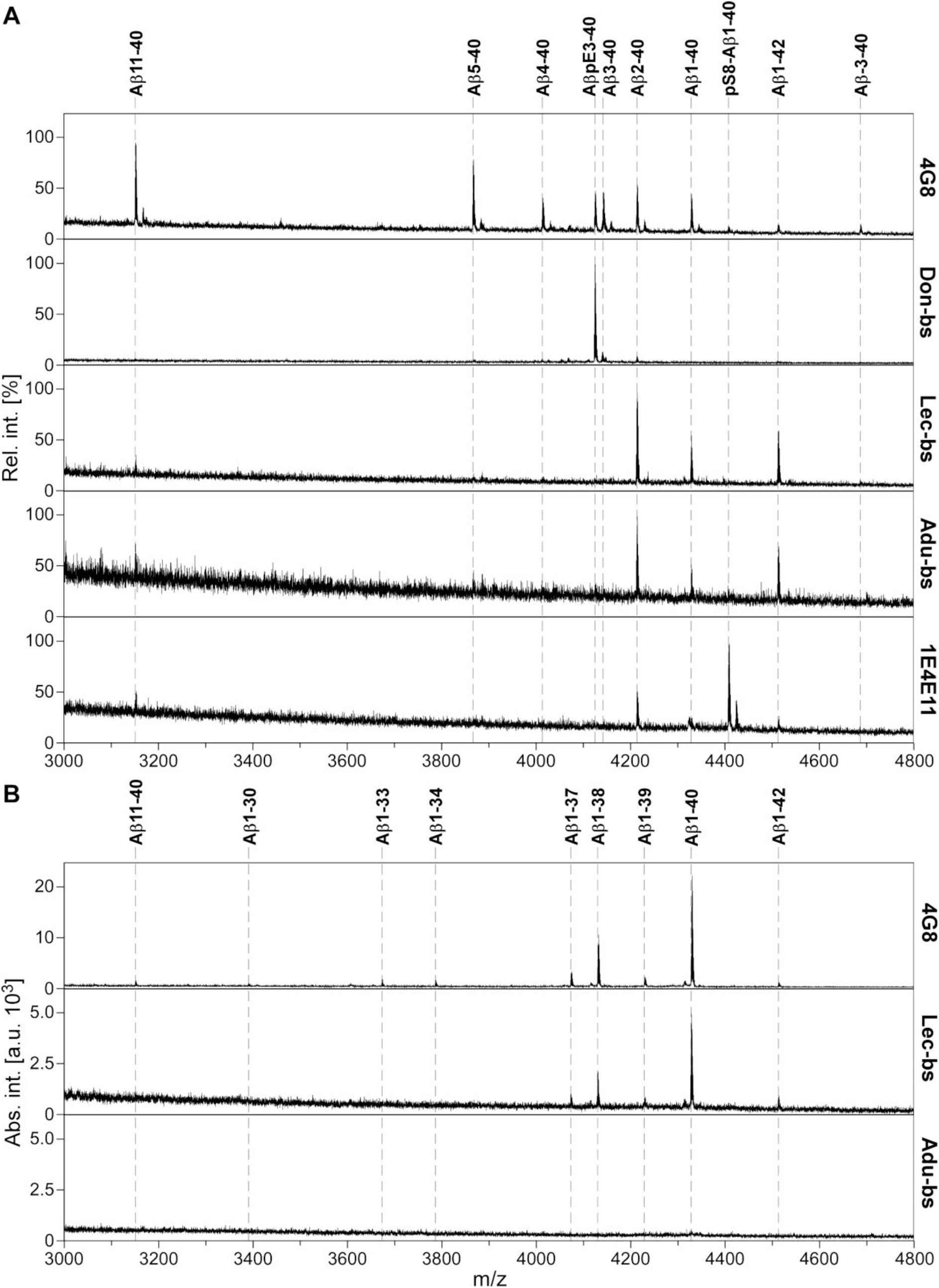
IP-MS signature of different biosimilar Aβ-Abs. **(A)** A mixture of synthetic Aβ peptides was immunoprecipitated with the following Aβ-Abs (from top to bottom): 4G8 (pan-antibody), Donanemab_bs_ (Don-bs), Lecanemab_bs_ (Lec-bs), Aducanumab_bs_ (Adu-bs), and 1E4E11 (phospho-selective control). Mass spectra were generated by MALDI-TOF-MS in reflector mode from TOPAC matrix and are representative for two independent experiments. We noted that the signal pattern slightly varied between experiments, which we mainly attributed to some immunoprecipitation of co-aggregated Aβ species, an effect that has to be considered under the experimental conditions applied. (B) IP-MS from pooled human CSF using MALDI-TOF-MS from conventional CHCA matrix. Mab 4G8 (positive control) and Lecanemab_bs_, but not Aducanumab_bs_ immunoprecipitated authentic biological Aβ peptides from pooled human CSF under the tested conditions. Major peaks were observed for Aβ1-40 and Aβ1-38. The spectra shown are representative for two replicate measurements each from two independent immunoprecipitations.

The IP-MS signature of 4G8, after incubation with the 10 peptide mix, showed signals from all constituents thus acting as positive control, with major intensities for Aβ5-40, intermediate signals for Aβ4-40, Aβ3-40, Aβ2-40, and Aβ1-40, and only minor signals for Aβ1-42, Aβ(-3- 40) and pSer8 Aβ1-40 (**Figure 4A**). Additionally, pyroglutamate AβpE3-40 was clearly recognized by mAb 4G8. As expected from its central epitope 17-21, 4G8 was the only Ab that showed an intense signal for N-terminally truncated Aβ11-40. The monoclonal antibody 1E4E11 preferentially precipitated pSer8-Aβ1-40-amide and, additionally, Aβ2-40, Aβ1-42 and Aβ1-40. The phosphorylated pSer8-Aβ1-40-amide produced the highest signal, suggesting a certain degree of preference (but not specificity) of 1E4E11 for phosphorylated pSer8-Aβ- amide under the tested conditions (**Figure 4A**).

In line with the PTM-microarray screening (**Figure 3**), Donanemab_bs_ exhibited highly preferential binding to AβpE3-40, with only minor signals for Aβ3-40 and Aβ2-40 (**Figure 4A**). The IP-MS spectra obtained with Lecanemab_bs_ showed predominant peaks for Aβ2-40, Aβ1-42, and Aβ1-40 in line with the apparent location of the epitope in the N-terminal region of Aβ (**Figure 4A**). However, synthetic Aβ peptides may display different properties than authentic Aβ peptides in biological samples, such as human cerebrospinal fluid (CSF). Importantly, Lecanemab was previously shown to have a strong preference for Aβ protofibrils and to bind Aβ monomers only weakly [83]. To test whether Lecanemab_bs_ was able to immunoprecipitate endogenous soluble Aβ from a relevant biological sample, we repeated the Lecanemab_bs_ IP-MS experiment with 250 µL of pooled CSF. In the resulting IP-MS spectrum, Aβ1-40, Aβ1-38, Aβ42, Aβ1-37 and Aβ1-39 were detected, albeit with relatively low signal intensities (**Figure 4B**). IP-MS from pooled CSF with Aducanumab_bs_ did not produce any Aβ signals (**Figure 4B**). In contrast, using the mix of ten synthetic Aβ peptides for Aducanumab_bs_ magnetic bead IP, we observed clear signals for Aβ2-40, Aβ1-42, Aβ1-40, but not for phosphorylated pSer8-Aβ1-40-amide (**Figure 4A).** This was in contrast to the binding data from array and BLI towards the minimal monovalent epitopes (**Figure 3 – figure supplement 2 and 3).**

For Aducanumab_bs_, microarray, BLI and IP-MS data suggest three binding modes. One presumably aggregation specific site (*EFRHDS*) as predicted by previous molecular dynamics studies [84], primarily via neutral V_L_, where the Ser(8) would only contact the Tyr-92 [84], in line with previous data [41]. A second binding site (EFRHDpS) relevant for the binding of short synthetic N-terminal Aβ fragments as resolved by our pSer(8) MD primarily contacting the positively charged V_H_ chain paratope via Arg-317-318 and in agreement with the sulfate ion-containing structure (**Figure 3 – figure supplement 4)**. Lastly, in a similar way the binding of Aducanumab to monomeric A*EFRHDS* (Aβ2-8) can benefit from the presence of negatively charged sulfate ions [41] (pdb: 6CO3). Of note, a recent molecular dynamics study suggested the presence of an N-terminal neutral domain in Aβ protofibrils (Aβ:Fab contact Ser8:Tyr92) [34], which may contribute for the preferential binding of Aducanumab to aggregated forms of Aβ.

### Aggregation-Dependent Binding of Biosimilar Aβ-Abs in vitro and in AD brain tissue

To further complement the characterization of the biosimilar Aβ-Abs, we applied gel electrophoretic separations of Aβ peptides followed by Western blotting, automated Capillary Isoelectric Focusing immunoassays (CIEF-immunoassay) with charge-based separation of Aβ peptides, and immunohistochemistry (IHC) experiments on brain tissue with pre-adsorption of Aβ-Abs.

Aducanumab and Lecanemab preferably bind specific aggregated forms of Aβ over monomeric forms [38,40,41][38,83]. In agreement with these reports, our analysis showed that Aducanumab_bs_ clearly detected highly aggregated fibrillar Aβ1-42 on Bistris-SDS gradient PAGE/Western blot. In contrast, it generated essentially no appreciable signals with monomerized hexafluoroisopropanol (HFIP)-treated or oligomeric Aβ1-42 preparations under the tested conditions (**Figure 5A**). On peptide Bicine-Tris SDS-PAGE/Western blot and on CIEF-immunoassays we observed substantial variances with Aducanumab_bs_ between different experiments (data not shown), making reliable interpretations difficult [85]. The observed high variance of the Aducanumab_bs_ results on CIEF-immunoassay and peptide SDS-PAGE/Western blot might be related to the low affinity of Aducanumab for monomeric Aβ [41] and the possibility that Aβ dimers or low order oligomers might form during the experiment [85]. In good agreement with the strong preference of Aducanumab_bs_ for highly aggregated fibrillar Aβ (see above), pre-adsorption with fibrillar Aβ1-42 largely blocked Aducanumab_bs_ immunoreactivity of vascular and parenchymal Aβ deposits in AD brain tissue, in contrast to pre-adsorption with an excess of Aβ1-42 monomers (**Figure 5B**).

**Figure 5:**
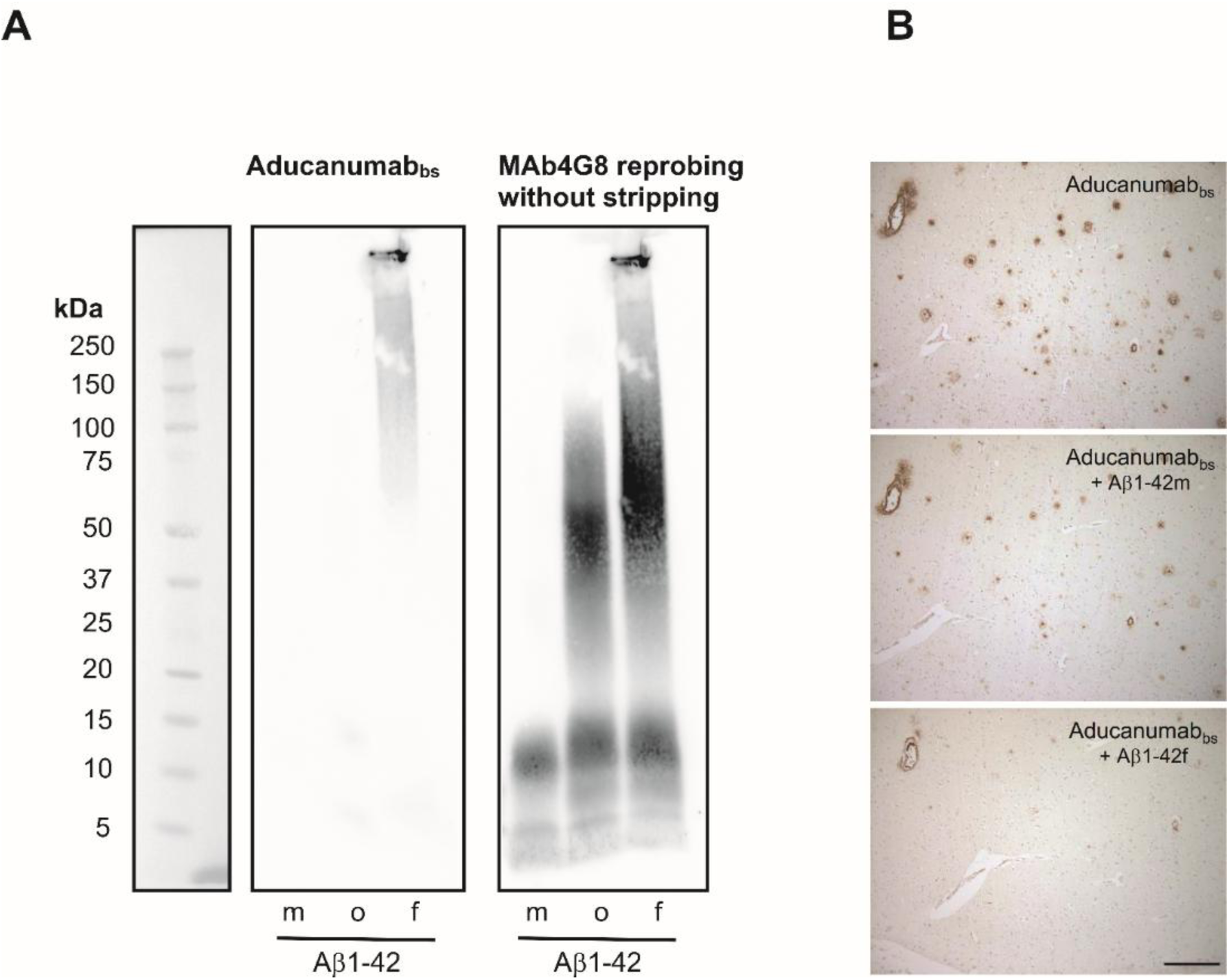
Aducanumab_bs_ shows strong preference for highly aggregated fibrillar Aβ1-42. (A) HFIP-treated Aβ1-42 monomer (m), oligomer (o) and fibril preparations (f) (1.44 µg each) were separated by 4-12% Bistris SDS-PAGE, blotted onto nitrocellulose and probed with Aducanumab_bs_ (5 min exposure, F 0.84, image display: High: 65535; Low: 0; Gamma: 0.85). Following image recording the blot membrane was reprobed without prior stripping (no removal of primary and secondary antibodies) with mAb4G8 (1 min exposure, image display: High: 54971; Low: 0; Gamma: 0.99). That way the 4G8 signals were essentially added on top of the initial Aducanumab_bs_ signals and background artifacts. The positions of pre-stained protein marker bands on the blot membrane are shown on the left-hand side. (B) Parallel sections from an AD patient were stained with Aducanumab_bs_ and Aducanumab_bs_ that had been pre-adsorbed with either Aβ1-42 monomers (Aβ1-42m) or Aβ1-42 fibrils (Aβ1-42f). Scale bar: 200 µm.

Lecanemab_bs_ recognized Aβ1-40, Aβ1-42, Aβ3-40 (pI approximately 5.97) Aβ-3-40 (aka. APP669-711),phosphorylated pSer8-Aβ1-40on CIEF-immunoassay and Bicine-Tris SDS-PAGE/ Western blot, thus recapitulating the initial microarray mapping for Lecanemab_bs_ shown in **Figure 1**. It appears, that under these experimental conditions Lecanemab_bs_ can bind to an epitope located in the N-terminal region of the Aβ sequence. On gradient Bistris SDS-PAGE/Western blot and under the tested conditions, Lecanemab_bs_ showed a certain, but not absolute preference for aggregated forms of Aβ1-42 over monomeric and low-n oligomeric forms in the HFIP-treated Aβ1-42 preparation (**Figure 6**). The immunohistochemical detection of Aβ in vascular and parenchymal deposits by Lecanemab_bs_ in brain sections from an AD- patient was efficiently blocked by antibody pre-adsorption with monomerized, oligomeric and fibrillar preparations of Aβ1-42 (**Figure 6E**). However, it cannot be excluded, that the HFIP treated Aβ1-42 monomers aggregated during the experiment.

**Figure 6:**
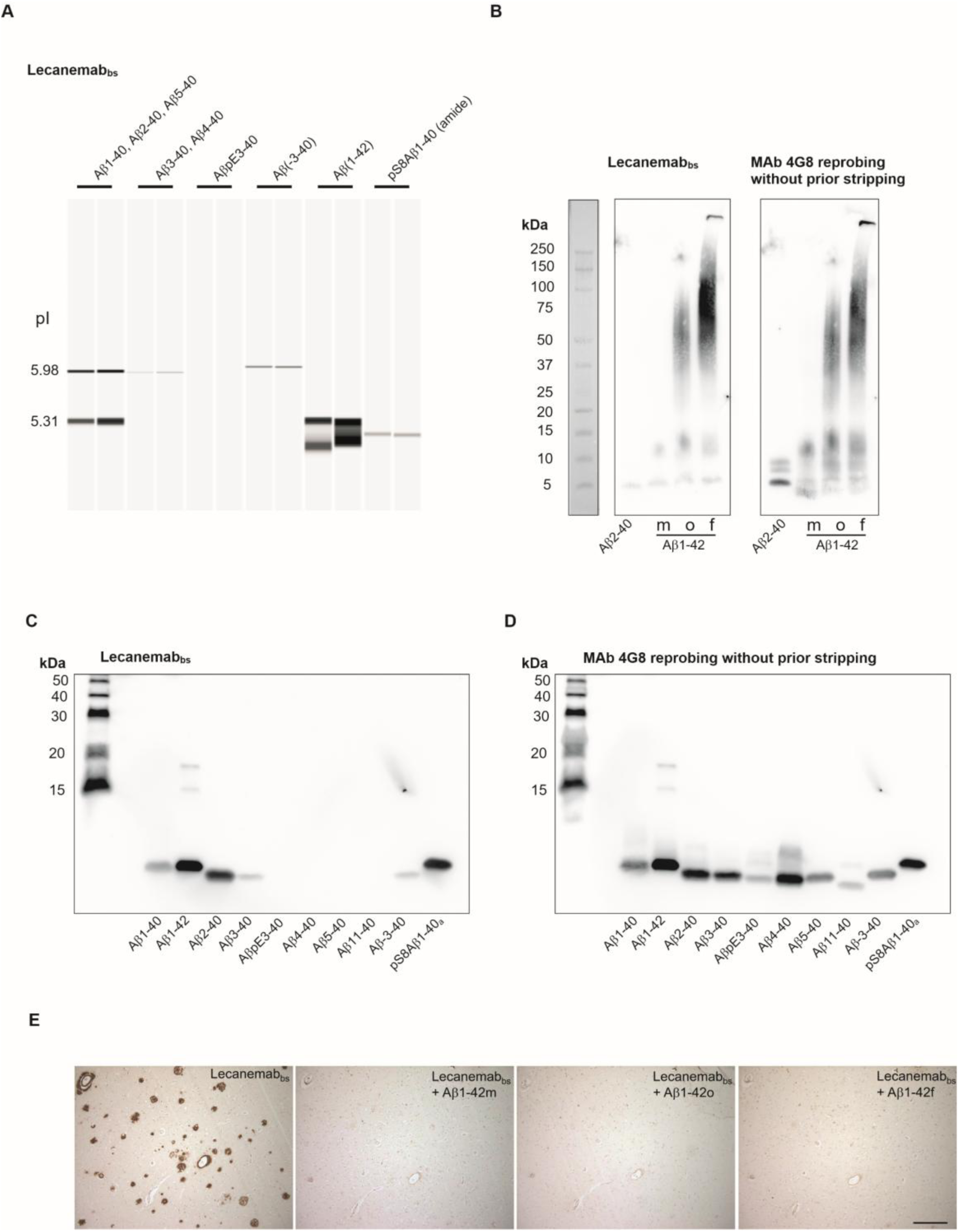
Lecanemab_bs_ recognizes an epitope located in the N-terminal region. (A) shows an example of CIEF-immunoassay results for Lecanemab_bs_ displayed as lane view (Western blot simulation). The indicated Aβ peptides were separated by isoelectric focusing in microcapillaries on a Peggy Sue device, immobilized photochemically to the inner capillary wall and subjected to immunological detection. Under the tested conditions, Lecanemab_bs_ detected Aβ1-40 (pI 5.31), Aβ2-40 (pI 5.98), Aβ3-40 (pI 5.97), Aβ-3-40 (pI 6.04), Aβ1-42 (pI approx. 5.3) and pSer8Aβ1-40 (amide) (pI 5.13-5.14). (B) Synthetic Aβ2-40 (361 ng), and monomerized (m), oligomeric (o) and fibrillar (f) preparations of Aβ1-42 (361 ng of each) were separated by 4-12% Bistris SDS-PAGE, blotted onto nitrocellulose and probed with Lecanemab_bs_ (0.5 µg/mL). Exposure: 2 min 7.5 sec; F 0.84; image display: high: 47545; low: 0; gamma: 1.0. After blot development, the blot membrane was reprobed with mAb 4G8 without prior stripping. That way the 4G8 signals were essentially added on top of the initial Lecanemab_bs_ signals and background artifacts. Exposure: 1 min F 0.84; image display: high: 46277; low: 0; gamma: 1.0. (C) The indicated Aβ peptides (50 ng of each) were separated by Bicine Tris (peptide) SDS-PAGE, blotted on PVDF, and probed with Lecanemab_bs_. Exposure: 5 min F 0.84, image display: high: 65535; low: 0; gamma: 0.85. (D) To confirm loading and successful blotting of all Aβ peptides, the blot membrane was re-probed with mAb 4G8 without prior stripping (see above). Exposure: 5 min F 0.84; image display: high: 65535; low: 0; gamma: 1.0. (E) Parallel sections from an AD patient were stained with Lecanemab_bs_ and Lecanemab_bs_ that had been pre-adsorbed with either Aβ1-42 monomers (Aβ1-42m), Aβ1-42 oligomers (Aβ1-42o) or Aβ1-42 fibrils (Aβ1-42f). Scale bar: 200 µm.

### C-Terminal Aβ Elements Contribute to Lecanemab-Binding in Tissue

To investigate in more detail the dependence of Lecanemab_bs_ binding on the Aβ peptide length and presumed resulting conformational differences, we performed additional pre-adsorption assays using a series of C-terminal truncated Aβ peptides with an intact N-terminal epitope. Since the original Lecanemab biosimilar antibody (Proteogenix Cat. No PX-TA1746) was not available anymore, Lecanemab_bs_ obtained from a different source (BioXCell (#BXC-SIM0032/ InVivoSIM) was used. Again, preincubation with Aβ1-42 monomers (Stressmarq) completely blocked Lecanemab_bs_ binding, whereas shorter peptides such as Aβ1-16, Aβ1-34 and Aβ1-38 did not. Thus, it appears that an intact N-terminus alone is not sufficient for Lecanemab_bs_ recognition, suggesting that additional elements contribute to Lecanemab binding of oligomers and aggregates.

On CIEF-immunoassay and Bicine-Tris SDS-PAGE/Western blot, Donanemab_bs_ detected AβpE3-40 and Aβ3-40 (both at a pI of approximately 5.95 on CIEF immunoassay) with a clear preference for the Aβ variant starting with the cyclized N-terminal pE. Other N-terminal Aβ variants that were tested produced no or very small signals (**Figure 7 B, C**). On gradient Bistris SDS-PAGE/Western Blot, Donanemab_bs_ detected multiple AβpE3-40 bands in the low Mwrange, presumably representing monomers and low order oligomers which were formed during sample preparation or SDS electrophoresis [85]. The HFIP-treated monomeric, oligomeric and fibrillar Aβ1-42 preparations produced very faint signals with Donanemab_bs_ (**Figure 7D**). Re-probing of the blot (without stripping) with mAb4G8 showed multiple Aβ1- 42 bands in the low molecular weight range (<15 kDa) in the HFIP-treated preparation (presumably monomers and low order oligomers). The Aβ1-42 oligomer preparation showed a similar pattern of bands smaller than 15 kDa and between 25 and 100 kDa. The Aβ1-42 fibril preparation showed low order oligomeric Aβ1-42 bands, in the molecular weight range of 37.5 to 150 kDa as well as highly aggregated Aβ1-42 which, unable to enter the stacking gel. On immunohistochemistry, the detection of Aβ vascular and parenchymal deposits by Donanemab_bs_ in AD brain sections was efficiently suppressed by pre-adsorption of the antibody with synthetic AβpE3-40, but not with Aβ1-40 or Aβ3-40 peptides (**Figure 7E**).

**Figure 7:**
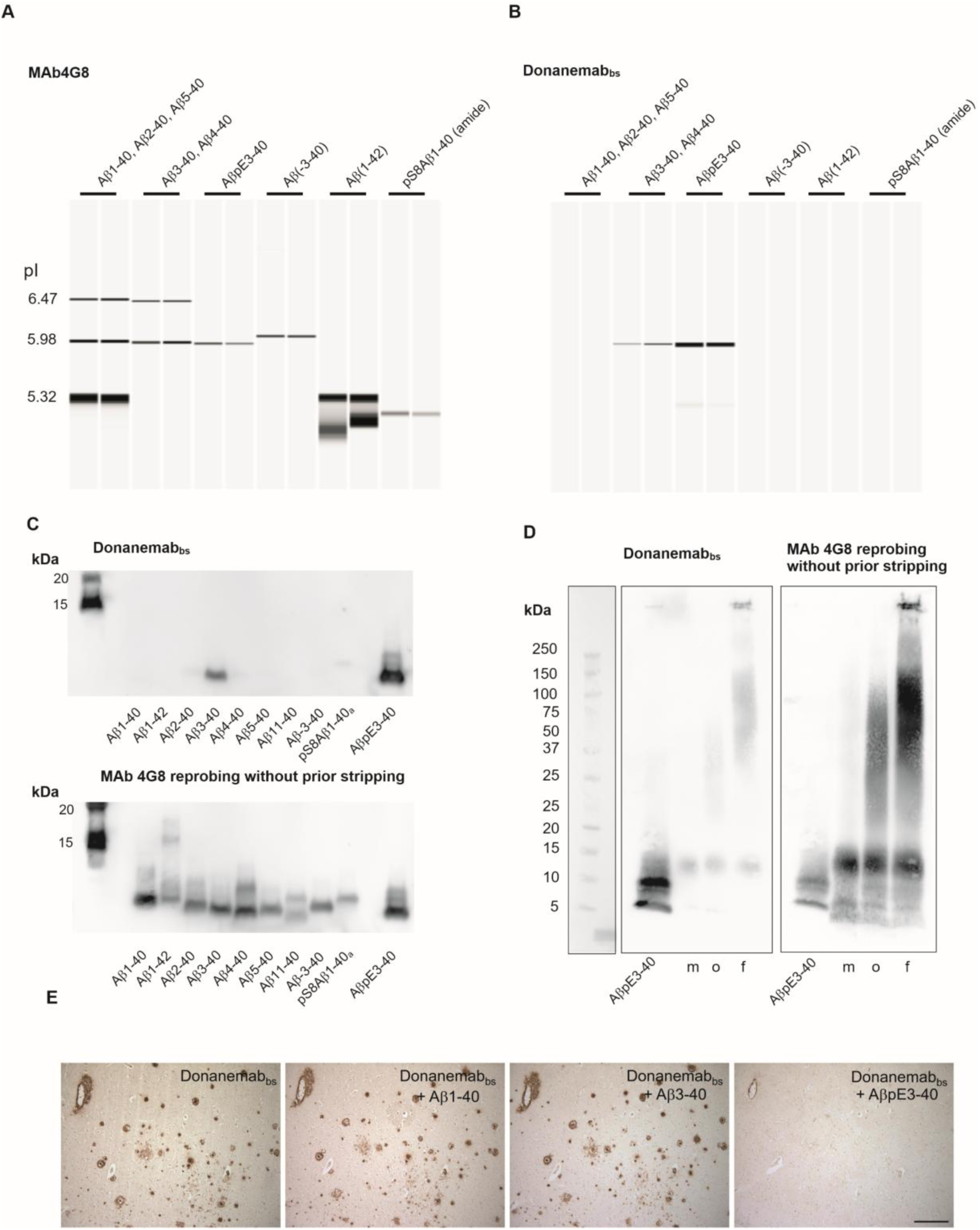
Donanemab_bs_ shows high preference for pyroglutamate bearing pEAβ3-40. The upper panel shows lane views (Western blots simulation) of a CIEF-immunoassay for assessing the detection of different Aβ variants by (A) mAb 4G8 (positive control antibody) and (B) Donanemab_bs_. The indicated Aβ variants were separated by isoelectric focusing in microcapillaries on a Peggy-Sue device, immobilized photochemically to the inner capillary wall and subjected to immunological detection by (A) mAb 4G8 and (B) Donanemab_bs_. (C) Synthetic Aβ peptides (50 ng per lane) were separated by Bicine Tris (peptide) SDS-PAGE, blotted on PVDF and probed with Donanemab_bs_ (0.5 µg/mL) (upper image). Exposure: 5 min F 0.84; image display: high: 59406; low: 0; Gamma: 0.65. To confirm that all Aβ variants were loaded and blotted, the blot membrane was reprobed with mAb 4G8 without prior stripping. That way, the 4G8 signals were essentially added on top of the initial Donanemab_bs_ signals and background artifacts (lower image). Exposure: 5 min F 0.84; image display: high: 65535; low: 0; gamma: 0.77. (D) AβpE3-40 and Aβ1-42 monomer (m), oligomer (o) and fibril preparations (f) were separated by 4-12% Bistris SDS-PAGE, blotted onto nitrocellulose and probed with Donanemab_bs_ (image display: High: 64091; Low: 0; Gamma: 0.75), followed by reprobing without prior stripping with mAb4G8 (image display: High: 46984; Low: 0; Gamma: 1.0). The positions of prestained protein marker bands on the blot membrane are shown on the left-hand side. (E) A pre-adsorption experiment employing synthetic Aβ peptides in human AD brain tissue showed that the immunoreactivity of Donanemab_bs_ was not suppressed by pre-adsorption of the antibody with excess amounts of synthetic Aβ1-40 and Aβ3-40 peptides. In contrast, a pre-incubation with AβpE3-40 effectively attenuated the antibody signal in extracellular Aβ deposits as well as in the vasculature.

## Conclusion

The exact binding specificity of Aβ-Abs, and its dependence on sequence context and aggregation state, is central both for their use as research tools and for their evaluation and development as therapeutic and diagnostic agents. Here we combined high-density peptide microarray analysis with complementary methodologies to generate a comprehensive dataset for 20 research and biosimilar Aβ-Abs. In total, 20,000 peptide–antibody interactions resolved individual epitopes at single–amino-acid resolution and clarifying the influence of PTMs and amino-acid exchanges for all epitopes within a unified experimental setup. The array analyses were complemented by IP-MS, CIEF immunoassay, SDS-PAGE/Western blot, and immunohistochemistry studies, thereby linking fine epitope definition and profiling with binding of full-length Aβ under native and aggregated conditions.

Comparative analyses highlighted clear mechanistic differences between the three therapeutic antibodies. For Donanemab, screening of monovalent peptides consistently recapitulated a core epitope at residues 3–10, with cyclized AβpE3-x as the dominant binding determinant. In contrast, Lecanemab and Aducanumab both required aggregated Aβ for full binding activity but mapped to overlapping N-terminal sequences (residues 3–7 for Lecanemab; 2–7 for Aducanumab). Binding quantification further substantiated a preference of Aducanumab for Ser8-phosphorylated Aβ for non-aggregated forms, and a marked sensitivity to the familial H6R mutation, consistent with recent reports linking antibody efficacy to differences between soluble and aggregated Aβ pools [86]

The qualitative and quantitative binding data, together with our molecular dynamics simulations and the previous structural reports, provide a mechanistic explanation for the observed binding requirements. For Aducanumab, X-ray crystallography revealed a Fc-mediated dimer interface that stabilises binding to aggregated Aβ [41]. This dimerisation also explains our observed preference for pSer8-modified Aβ monomers: phosphorylation neutralises the Arg-Ala-Arg motif at the Fc-dimer interface, thereby promoting dimerisation and avidity-enhanced binding. In aggregated Aβ forms, such phosphorylation is dispensable because avidity is already provided by multimerisation, reconciling our peptide library, BLI, and tissue binding results with structural models and simulations of Aducanumab [41, [84],93]. For Lecanemab, our binding profiles align closely with the model proposed on the basis of D3 antibody similarity (PDB 5MY4) [93], correctly accounting for sequence tolerances at positions 3–7. However, in contrast to published claims [93], we find no evidence that N-terminal extensions beyond Glu3 reduce affinity. Binding is preserved both in vitro and in AD tissue, supporting a minimal epitope definition that includes residues 3–7, consistent with our peptide, MS, and immunohistochemistry data [38,93], and with recent demonstrations of Lecanemab’s preferential binding to fibrillar Aβ aggregates [46].

In summary, we systematically compared the binding requirements of 20 Aβ-Abs, addressing sequence (including genetic and species variants), PTM, and structural aspects. By resolving discrepancies between structural models and biochemical data, we provide mechanistic insight into the distinct binding modes of Aducanumab and Lecanemab, and into the strict selectivity of Donanemab. This work provides a resource for the rational selection of Aβ-Abs in research and diagnostics and may help inform the evaluation and design of future therapeutic and diagnostic Aβ-Abs.

## Methods

### Ethics approval and patients consent

In this study, a pooled CSF sample prepared from several individual CSF samples from the local biobank at the Department of Psychiatry and Psychotherapy at the University Medical Center Göttingen was used in some IP-MS experiments. The collection and archiving of biological samples and clinical data in strictly pseudonymous form in a local biobank and their use in biomarker research were approved by the ethics committee of the University Goettingen (Protocol number 9/2/16). From all participants or their legal representatives written informed consent was obtained prior to the inclusion in the biobank. All study participants received detailed information prior to enrolment in the study and collection of the biomaterial. In addition, an information sheet was handed out by the study physicians at the Department of Psychiatry and Psychotherapy at the University Medical Center Göttingen. After a possible reflection period, written informed consent was obtained from the study participants. If the ability to consent was not given, a separate informed consent form was also provided for the study participant’s legal representative, and the legal representative signed the informed consent form for the study. All procedures involving human participants were in accordance with the ethical standards of the institutional and/or national research committee and with the 1964 Helsinki declaration and its later amendments or comparable ethical standards.

Paraffin-embedded human brain samples from individuals with sporadic Alzheimer’s disease (AD) were obtained from the Netherlands Brain Bank (NBB,Netherlands Institute for Neuroscience, Amsterdam (open access: www.brainbank.nl)). All material has been collected from donors for or from whom a written informed consent for a brain autopsy and the use of material and clinical information for research purposes had been obtained by the NBB. For the -analyses in the present study, sections from the medial frontal gyrus region of the brain were utilized. All procedures involving human tissue were approved by the ethical committee at the University Medical Center Göttingen (Protocol number 12/1/15).

### Preparation of Functionalized Cellulose Discs

Paper functionalization was performed by introducing some variations to the previous protocol [87] aimed at increasing the output of discs per session. In brief, Whatman paper grade 50 (45 × 25 cm) was cut into 4mm diameter discs using an OMTech 50-watt CO2 laser at 7.8% power and 150 mm/s speed. To reduce mechanical stress and facilitate transfer into a 384-plate, each disc was cut with a semi-circular pattern. The pre-cut layout (15 x 10 cm) was then treated for 3 hours with a solution of 9-fluorenylmethyloxycarbonyl-β-alanine (Fmoc-β-Ala-OH), diisopropylmethanediimine (DIC) and 1-Methylimidazole (NMI). The Fmoc absorption of each batch of functionalized cellulose membrane was determined at 290nm after 20 minutes of deprotection with 20% (v/v) piperidine (pip) in DMF. The average loading was 130nmol/disc.

### Microarray Synthesis

The µSPOT assay was performed as previously described [88]. First, the APP sequence (UniProtKB: P05067) was displayed in microarray format as 15mer overlapping peptide library. Peptide arrays were synthesized [89–91] using MultiPep RSi robot (CEM GmbH, Kamp-Lintford, Germany) on cellulose discs containing Fmoc-β-Ala linkers. Natural amino acid building blocks were purchased from IRIS (IRIS Biotech GmbH, Marktredwitz, Germany). Special building blocks employed in this study such as isomers, stereoisomers, phosphorylation and pyroglutamination post-transcriptional modifications are listed here: Fmoc-L-Ser(PO(OBzl)OH)-OH (Iris – FAA1423), L-Pyroglutamic acid pentachlorophenyl ester (Novabiochem - 8.54203), Fmoc-L-Asp-OtBu (Iris – FAA1356), Fmoc-D-Asp-OtBu (Carbosolutions – CC05095) and Fmoc-D-Asp(tBu)-OH (Iris – FAA1309). Synthesis was carried out by deprotecting the Fmoc-group using 20% piperidine in dimethylformamide (DMF). Peptide chains were elongated using a coupling solution consisting of amino acids (0.5M) with oxyma (1M) and DIC (1M) in DMF (1:1:1). Coupling steps were carried out 3 times (30min), followed by capping (4% acetic anhydride in DMF). Discs were transferred into 96 deep-well plates for the work-up. Side chains were deprotected using 90% trifluoracetic acid (TFA), 2% dichloromethane (DCM), 5% H_2_O and 3% triisopropylsilane (150μL/well) for 1h at room temperature (RT). Afterward, the deprotection solution was removed, and the discs were solubilized ON at RT, while shaking, using a solvation mixture containing 88.5% TFA, 4% trifluoromethanesulfonic acid (TFMSA), 5% H2O and 2.5% TIPS (250μL/well). The resulting peptide-cellulose conjugates (PCCs) were precipitated with ice-cold ether (700μL/well) and spun down at 2000×g for 10min at 4°C, followed by two additional washes of the formed pellet with ice-cold ether. The resulting pellets were dissolved in DMSO (250μL/well).

### Microarray Quality Control

LC-MS was carried out using peptide quality controls that were cleaved from solid support. To ensure cleavage, a Rink amide linker (Iris) suitable for SPPS on cellulose support was introduced during the first coupling cycle. In an acidic environment, the quality controls were cleaved off from the solid support. To isolate the quality controls, 150 µL of the supernatant (SN) was transferred to 1.5 mL reaction tubes, followed by the addition of 700 µL of diethyl ether. The samples were then vortexed, and the peptides were allowed to precipitate by incubation at -20°C overnight. After centrifugation at 13,300 × g and 4°C for 10 min, the SN was discarded, and 500 µL of diethyl ether was added. The mixture was vortexed, centrifuged for 10 min, and the supernatant was decanted. This process was repeated twice, and the peptides were left to dry for 60 min. Finally, the Rink amides were dissolved in 50 µL of 50% acetonitrile: 0.1% formic acid (v/v) and vortexed briefly before centrifugation at 13,300 × g and room temperature. For analysis, the quality controls were diluted 1:3 and sent to LC-MS (Agilent technologies).

### Microarray Printing

PCCs solutions were mixed 2:1 with saline–sodium citrate buffer (150mM NaCl, 15mM trisodium citrate, pH7.0) and transferred to a 384-well plate. For transfer of the PCC solutions to white-coated CelluSpot blank slides (76×26mm, Intavis AG Peptide Services GmbH and CO. KG), a SlideSpotter (CEM GmbH) was used. After completion of the printing procedure, slides were left to dry ON.

### Microarray Binding Assay

The microarray slides were blocked for 60min in 5% (w/v) skimmed milk powder (Carl Roth) 0.05% Tween20 phosphate-buffered saline (PBS; 137mM NaCl, 2.7mM KCl, 10mM Na_2_HPO_4_, 1.8mM KH_2_PO_4_, pH7.4). After blocking, the slides were incubated for 30 min with a solution of blocking buffer and monoclonal and polyclonal Aβ-Ab (**Table 1**). They were then washed 3× with PBS 0.05% Tween20 for 1min. IgG antibodies were detected using goat anti mouse IgG-HRP (Thermo Fisher Cat. No31430), Goat anti-Rabbit IgG, Antibody, HRP (Cell Signaling: 7074), Goat anti-Rat IgG (H+L) Secondary Antibody, HRP (TF: 31470) and Goat anti-Human IgG (H+L) Secondary Antibody, HRP (TF: 31410). The chemiluminescence readout was detected with an Azure imaging system c400 (lowest sensitivity, 1s, 5s, 10s, 30s, 1m, 2m) using SuperSignal West Femto maximum sensitive substrate (Thermo Scientific GmbH, Schwerte, Germany). The raw grayscale intensities were measured using open-source MARTin software [92].

### Dot Blot

The purified peptides were printed onto white-coated CelluSpot blank slides using the Slide Spotter at a starting amount of 0.05 µg. The slides were allowed to dry for 3 hours and rinsed once with 1X PBS to remove the unbound peptides. After blocking in 5% (w/v) skimmed milk powder for 1 hour, the slides were incubated with 1nM Aducanumab_bs_ and IgG signals were detected using goat anti-human IgG DL650 (1:1000 - Invitrogen 84545). The fluorescence signal readouts were acquired using Amersham Image Quant 800 (GE Healthcare) with an exposure time of 30 seconds and 4x4 binning.

### Molecular Dynamics Simulations

For molecular dynamic (MD) simulations the crystal structure of the aducanumab Aβ complex structure (PDB: 6CO3) was used. The complex structure was prepared using MOE (Molecular Operating Environment, 2022.02, Chemical Computing Group ULC, 1010 Sherbooke St. West, Suite #910, Montreal, QC, Canada, H3A2R7, 2023) structure preparation tools with default values. For producing the MD-starting structure the experimentally resolved residues of the Aβ were prolonged by either a serine or a phosphorylated serine in MOE.

The MD simulations were conducted using AMBER22 suite [93]. Topology and parameter files were generated in tleap module using the AMBER ff14sb. The simulations were set up in the following way: The peptide-aducanumab complex was energy minimized for 2000 steps using a generalized Born implicit solvent model, neutralized, and solvated using the TIP3P water model. The system was equilibrated by heating the solvent from 100 K to 300 K within 500 ps under constant volume conditions with non-solvent atoms assigned with harmonic constraints of 100 kcal·mol-1·Å-2. Cooling the system back to 100 K within 250 ps and lowering the harmonic constraints on non-solvent atoms to 1 kcal·mol-1·Å-2, followed by heating to 300 K within 500 ps period. Afterwards, the non-solvent atoms constraints were set off and the pressure was adjusted to 1 bar using the Berendsen barostat. Each complex (unmodified and phosphorylated) was simulated in triplicates for 200 ns at constant pressure and temperature. All simulations were carried out using periodic boundary and particle mesh Ewald (PME) methodology to handle long-range electrostatics. 2 fs-time steps were used for integration and coordinates of were saved every 500 steps (1 ps). The trajectories were further analysed with CPPTRAJ (4.14.0).

### Aβ-Peptides for SDS-PAGE, CIEF Immunoassay and IP-MS

The synthetic peptides Aβ1–40, Aβ1–42, Aβ2–40, Aβ3–40, AβpE3–40, Aβ4–40, Aβ5–40 and Aβ11–40 were obtained from AnaSpec (Fremont, California, USA 94555). . Synthetic Aβ−3– 40 [94]was kindly provided by H.-J. Knölker, Technische Universität Dresden. Phosphorylated pSer8-Aβ1-40 was synthesized with an amidated C-terminal carboxyl group to remove one negative charge, ensuring that this Aβ variant still falls into the applied separation range of the CIEF immunoassay.

### Bistris-SDS Gradient PAGE/Western Blots Analysis of Different **Aβ Aggregation States**

For the assessment of antibody binding to different Aβ aggregation states, we used Aβ1-42 monomers (Cat. No SPR-485B), Aβ1-42 oligomers (Cat. No SPR-488B) and preformed Aβ1- 42 fibrils (Cat. No SPR-487B) (StressMarq biosciences, Victoria, BC, Canada). The samples for electrophoresis were prepared (starting from stock solutions) with final concentrations of 0.1 µg/µL) in LDS sample buffer (247.5 mM Tris-HCl, pH 8.5, 2% (w/v) lithium dodecyl sulfate (LDS), 10% (v/v) glycerol, 0.5 mM EDTA, 0.22 mM Brilliant Blue G250, 0.175 mM Phenol Red) and denatured by heating at 70°C for 10 min. Per lane, 0.361, 0.722 or 1.44 µg of the different Aβ1-42 forms were loaded on a 4-12% Bistris-gel (1mm, 12 wells, Anamed Electrophorese GmbH, Groß-Bieberau / Rodau, Germany) and separated at 200 V using MES-SDS running buffer. Following recommendations provided by the manufacturer of the different Aβ1-42 aggregation forms, electroblotting onto nitrocellulose was done at 100 V for 1 hour with a Biorad mini tank blot. The prechilled transfer buffer contained 25 mM Tris, 192 mM glycine, 20% (v/v) methanol and 0.0375 % Tween-20. After blotting, the membrane was washed for 10 min at room temperature in Tris-buffered saline (TBS) followed by blocking for 30 min with 5% powdered milk in TBS containing 0.1% Tween-20 (TBS-T). The blots were then incubated with primary antibodies in 5% milk in TBS-T over night at 4°C and washed 3 x for 10 min with TBS-T at room temperature. The secondary antibodies were (i) Peroxidase-conjugated Affini Pure Donkey Anti-Human IgG (H+L), Jackons ImmunoResearch Laboratories Inc., West Grove, PA, (code number 709-035-149)1:20000 in TBS-T for Donanemab_bs_, Lecanemab_bs_ and Aducanumab_bs_ and (ii) anti-mouse IgG-POD (Calbiochem/EMD Millipore, San Diego, California, USA, catalog number 401253-2ML) 1:10000 in TBS-T for mAb 1E4E11 and mAb 6E10. After 1 h incubation with the secondary antibodies, the blots were washed 3 x for 10 min with TBS-T and incubated for 5 min with ECL-Prime Western Blotting Detection Reagent (Cytiva Amersham, Little Chalfont, England). Images were recorded on a Vilber-Loumart Fusion SL blot imager. To confirm successful blotting of all Aβ variants, the blot membranes were reprobed without prior stripping with mAb4G8 (0.5 µg/mL in 5% milk in TBS-T over night at 4°C or 1-2 h at room temperature) followed by anti-mouse IgG-POD (1:10000 in TBS-T) for 1 h at room temperature. That way, the 4G8 reprobing signals were essentially added on top of the initial signals and background artifacts.

### Peptide Bicine-Tris SDS-PAGE / Western Blot

Synthetic Aβ peptides were separated by 15% T/5% C Bicine-Tris SDS-PAGE, essentially according to[95]. After the electrophoresis, the gel was shortly rinsed with 50 mM Tris –HCl, pH 8.0, followed by blotting the peptides on PVDF using a Gravity Blotter (Serva Electrophoresis GmbH, Heidelberg, Germany). The blotting sandwich was assembled on the Gravity blotter from bottom to top by first placing the gel on the lower plate, followed by a precut PVDF membrane (activated with methanol and equilibrated in 50 mM Tris-HCl, pH 8.0), one filter paper Whatmann 3mM equilibrated in 50 mM Tris-HCl, pH 8.0, three dry filter papers (Sartorius BF4 blotting paper) and, finally, three 2.25 kg Gravity Blotter metal plates. The gravity/press blotting was done overnight at room temperature. On the next morning, the blot sandwich was disassembled and the PVDF membrane was boiled in PBS for 3min in a microwave oven with the protein side facing downwards. After blocking for 30 min at room temperature with 2% ECL Prime blocking reagent (Cytiva) in PBS containing 0.075% Tween- 20 (PBS-T), the blots were incubated for 1 h at room temperature with the primary antibodies (0.5 µg/mL in 2% ECL-Prime Western Blotting Detection Reagent in PBS-T). After 3 x 10 min washing with PBS-T, the secondary peroxidase-conjugated Affini Pure Donkey Anti-Human IgG (see above) (1:20000 in PBS-T) was applied for 1-2 hours at room temperature. Following 3 x 10 min washing with PBS-T, the blots were incubated for 5 min with ECL-Prime Western Blot Detection Reagent, and images were recorded. To confirm successful blotting of all Aβ variants, the blot membranes were reprobed without prior stripping with mAb4G8 (0.5 µg/mL in 2% ECL Prime blocking reagent in PBS-T over night at 4° C or 1 h at room temperature followed by anti-mouse IgG-POD (1:10000 in PBS-T) for 1 h at room temperature. That way, the 4G8 reprobing signals were essentially added on top of the initial signals and background artifacts.

### Capillary Isoelectric Focusing Immunoassay

Automated capillary isoelectric focusing immunoassays were performed with a Peggy-Sue system (ProteinSimple,San Jose, California, USA), essentially as previously described [96] and [53].

In brief, synthetic Aβ peptide variants were separated by isoelectric focusing in microcapillaries followed by photochemical immobilization to the inner capillary walls and probing with the primary antibodies for 120 min. After 2 washes, the horseradish peroxidase conjugated secondary antibodies were loaded and incubated for 60 min. After two washing steps, a luminol/peroxide mix was loaded, and chemiluminescent signals were recorded. The data was analyzed with the Compass for SW software. The following secondary antibodies were used: Anti-human IgG Secondary HRP antibody (Protein Simple, catalog number 043- 491), Goat-Anti-Mouse-HRP Secondary Antibody ProteinSimple, catalog number 040-655) and Peroxidase-conjugated Affini Pure Donkey Anti-Human IgG (H+L) (1:100 dilution) (Jackson ImmunoResearch Laboratories Inc., West Grove, PA, code number 709-035-149).

### Automated Solid Phase Peptide Synthesis

The peptides were synthesized using PurePep Chorus (Gyros Protein Technologies, US) with Fmoc chemistry at 0.05mmol scale. Tentagel S RAM (Resin Substitution 0.22mmol/g - Rapp Polymere, Germany) resin was first swollen in dimethylformamide (DMF) for 30-60 minutes. Next, Fmoc was removed using a 20% piperidine in DMF solution for 1-2 minutes and washed 3x with DMF. Deprotection steps were performed at 90°C for unphosphorylated peptides and at room temperature from the first phospho-serine coupling. The peptide chain was elongated by adding the premixed amino acid (AA, 125mM and Oxyma 125mM) along with N,N’- diisopropylcarbodiimide (DIC, 250mM). The mixture was heated at 90°C for 2 minutes and then washed 2x with DMF. Capping was done with acetic anhydride (10% in DMF) for 5 minutes and washed 3x with DMF. The peptide chain was elongated until completion and dried for 30 minutes after 7x DCM washes. The Fmoc deprotected peptides were cleaved from the resin using a mixture of 90% TFA, 5% water, 3% TIPS, and 2% DCM for 4 h at RT. The peptides were then precipitated in ice-cold ether ON, subsequently purified using HPLC and analysed by LC-MS, as described below.

### Purification and Characterization of Peptides

The crude peptides were purified by reverse phase (RP) HPLC using a water-acetonitrile gradient with 0.1% formic acid (FA). Preparative HPLC was performed on a UltiMate™ 3000 HPLC Systems instrument (ThermoFisher Scientific, Waltham, Massachusetts, USA) equipped with RS variable wavelength detector (set to 215nm) and a Kinetex® 5µm EVO C18 100 LC column (00D-4633-P0-AX; Phenomenex, Aschaffenburg, Germany). Chromatograms were recorded using Chromeleon software v7.2. Purity and structural identity of the peptides were verified using a DAD equipped 1260 Infinity II HPLC device with a C18 RP column (Onyx Monolithic C18 50×2 mm), coupled to a mass selective detector with a single quadrupole system (Agilent Technologies, Santa Clara, CA, US) in ESI+ mode.

Crude phosphorylated pSer8-Aβ1-40 with an amidated C-terminal carboxyl group was purified on a 10x150mm Waters (Milford, Massachusetts, USA) XBridge Protein BEH C4 OBD preparative column (300Å, 5 µm) using 3% acetonitrile/0.2% Formic acid as solvent A and 97% acetonitrile/0.2% FA as solvent B. A Beckman System Gold HPLC system with external column oven was used to form the following gradient with a flow rate of 3.8 ml/min at 50°C column temperature: 5-20% B in 2 min, 20-32% B in 22 min, 32-50% in 5 min. Analytical HPLC was performed on a Waters Alliance e2695 HPLC system coupled to a photodiode array (PDA) optic detector and a single quadrupole QDa mass detector. A 4.6x150mm Waters XBridge Protein BEH C4 analytical column (300 Å, 3.5µm) was used at a flow rate of 3.8ml/min at 50°C column temperature; the corresponding gradient was 5-20% B in 2 min, 20- 32% B in 11 min, 32-50% in 5 min.

### Biolayer Interferometry

The samples were dispensed into a 96 microplate (Greiner bio-one GmbH, Frickenhausen, Germany) at a final volume of 200 μL. The binding of Aducanumab biosimilar and mAb 1E4E11 and their phosphorylated, unphosphorylated and H6R epitopes 6xHis-tagged were measured using Biolayer interferometry on an Octet RED 96system (Sartorius, Göttingen, Germany). The peptides were synthesized (purity above 98% - based on UV at 215nm) with Fmoc-Adoa linker (Iris - FAA1435) and C-terminal 6xHis tag. Sensorgrams were measured in double reference against the buffer and the reference sensor signals.

NTA biosensors (Sartorius, Göttingen, Germany) were preincubated into 1XPBS buffer pH 7.4, 0.2% BSA, 0.02% (v/v) Tween 20 for 10 min. A first baseline was established in the same buffer. The 6xHis peptides were loaded onto NTA biosensors at 20 μg/ml for 200 s. Following a buffer wash of 120s, the biosensors were incubated with serial-dilutions of the respective antibodies for 1200s s to measure association and then into buffer for additional 1500s to observe the dissociation phase. Experiments were performed at 30 °C and sample plates were shaken at 1000 rpm. Sensorgrams were globally fitted using Data Analysis software version 12.0.2.3 (Sartorius, Göttingen, Germany) with a 1:1 binding model.

### Immunoprecipitation and Mass Spectrometry

Antibodies were covalently coupled to Dynabeads M-270 Epoxy (Invitrogen/ThermoFisher Scientific, Waltham, Massachusetts, USA Cat. No. 14311 D), essentially following the manufacturer’s instructions. Tween-20 was omitted from all wash and storage buffers for specific use in IP-MS. In brief, 5 mg of M-270 epoxy beads were weighed into a 1.5-mL reaction vial and washed with 1 mL of buffer C1 (kit component). The bead suspension was divided in aliquots of 200 µL (containing 1mg of beads). The vials were placed on a magnetic stand for a few minutes, and the supernatant was carefully removed and discarded. Washed beads were resuspended in 40 µL of buffer C1 and 10 μL of antibody stock solution (1 mg/mL). After gentle mixing by pipetting, 50 μL of buffer C2 (kit component) was added and incubated overnight at 37 °C on a mixer. After coupling, the beads were washed one time with 400 μL of buffer HB (kit component), one time with 400 μL of buffer LB (kit component) and two times with 400 μL of PBS. After a final wash with PBS for 15 min with continuous agitation, the beads were resuspended in 100 µL PBS/0.1% BSA/0.02% NaN_3_ and stored at 4 °C.

The synthetic Aβ peptides Aβ1–40, pSer8-Aβ1–40, Aβ1–42, Aβ2–40, Aβ3–40, AβpyGlu–40, Aβ4–40, Aβ5–40 and Aβ11–40 Aβ−3–40 (see above), were mixed and diluted to final concentrations of 10ng/mL of each in PBS/0.1%BSA containing phosphatase inhibitors (PhosphoStop Mini, Roche, Ref 04906837001) and protease inhibitors (Complete Mini Roche, Ref 04693124001). For immunoprecipitation of Aβ peptides, 250 µL of the Aβ mix or of pooled CSF was combined with an equal volume of 100 mM Tris-HCl, pH 7.4, containing 0.2% (w/v) nonyl-β-d-thiomaltoside (NTM) and 0.2% (w/v) n-dodecyl-β-d-maltoside (DDM) and 15 μL of functionalized M270 epoxy beads. After overnight incubation at 4 °C with constant mixing, the beads were washed five times with 0.5 mL of 50 mM Tris-HCl, pH 7.4, 150 mM NaCl, 0.1% (w/v) DDM, 0.1% (w/v) NTM and resuspended in 0.5 mL of 50 mM ammonium acetate (pH approximately 7.0). The beads were immediately forwarded to sample preparation for mass spectrometry. There, the beads were immobilised on a magnet, the supernatant was removed, and the beads were washed one time with 0.5 mL of 50 mM ammonium acetate, and one time with 0.5 mL H_2_O. The bound Aβ peptides were eluted in 2.5 μL of 70% acetonitrile containing 5 mM HCl.

Analysis by MALDI-TOF-MS was essentially performed as described [21,97]. Briefly, the eluates (0.5 µL) from synthetic samples were spotted onto a pre-structured MALDI sample support (MTP AnchorChip 384 BC; Cat. No. 8280790, Bruker, Bremen, Germany), followed by the addition of 0.5 µl TOPAC matrix solution, a matrix recently developed for the improved detection of intact phosphorylated Aβ peptides [82]. Alternatively, conventional CHCA matrix was used for CSF samples as described in [21]. A 1:1 mixture of Peptide Calibration Standard II (Cat. No. 8222570, Bruker, Bremen, Germany) and PepMix2 (Cat. No. C102, LaserBio Labs, Valbonne, France) was used as calibrant. The samples were dried in a vacuum desiccator and a total of 5000 mass spectra per spot were recorded using an ultrafleXtreme MALDI-TOF/TOF mass spectrometer in reflector mode operated under the software flexControl 3.4 (Bruker, Bremen, Germany).

### Immunohistochemistry and Antibody Preadsorption

Immunohistochemistry was performed on 4-μm paraffin sections. In brief, brain sections were deparaffinized in Roticlear (Carl Roth, Karlsruhe, Germany) and rehydrated in a descending series of ethanol. Activity of endogenous peroxidases was blocked by incubation in 0.3% H_2_O_2_ in PBS for 30 min. Antigen retrieval was achieved by initial boiling of the sections in 0.01 M citrate buffer (pH 6.0) and subsequent incubation in 88% formic acid for 3 min. Unspecific binding sites were blocked by incubation in PBS containing 4% skim milk and 10% fetal calf serum (FCS) for 1 h at room temperature, prior to the overnight incubation with the primary antibodies. The following antibodies were used for the detection of different Aβ variants: Aducanumab_bs_ (0.5 µg/µl), Donanemab_bs_ (1 µg/ml), Lecanemab_bs_ (0.06 µg/ml).

Antibody pre-adsorption with synthetic Aβ peptides (Aβ1-16 ( synthesized at Neuroproteomics Group, Max-Planck Institute for Multidisciplary Sciences (MPI-NAT), Göttingen), Aβ1-34, Aβ1-38, Aβ1-40, Aβ3-40 (from Anaspec), AβpE-40 (Peptide Speciality Laboratory, Heidelberg, Germany), Aβ1-42 monomers, Aβ1-42 oligomers and Aβ1-42 fibrils (StressMarq, Victoria, Canada) was carried out as described previously [52]. In brief, 1 µg Donanemab_bs_ was blocked with 2.5 µg of Aβ1-40, Aβ3-40 or AβpE3-40 in 0.01 M PBS including 10% FCS. In the case of Aducanumab_bs_, 0.5 µg of the antibody were blocked with 10 µg of Aβ1-42 monomers or 5 µg of Aβ1-42 fibrils. For Lecanemab_bs_, 0.06 µg of antibody (PX-TA1746) was blocked with 0.25 µg of Aβ1-42 monomers, oligomers or fibrils, or 0.06 µg (BXC-SIM0032) was blocked with 0.25 µg Aβ1-16, Aβ1-34, Aβ1-38 or Aβ1-42 monomers). Antibodies and peptides were incubated on a rotator for 3 h at room temperature and reaction tubes were centrifuged at 13000 rpm for 5 min at the end of the incubation period. Control conditions included antibodies in 0.01 M PBS including 10% FCS without synthetic peptides. Adjacent paraffin-embedded sections from a human AD case were incubated overnight with the supernatants and sections were washed with 0.01 M PBS including 0.1 % Tergitol. For detection, biotinylated anti-human antibodies (6 µg/ml, Jackson ImmunoResearch, West Grove, Pennsylvania, USA #109-065-003) were used and staining was visualized with the ABC method with a Vectastain kit (Vector Laboratories, Newark, California, USA), diaminobenzidine as a chromogen and counterstaining with hematoxylin.

## Author contributions

The following section was based on the CRediT system. Conceptualization: IT, HMM; Methodology: IT, HMM, HWK, OJ, OW; Formal Analysis: IT, HMM, TLe, HWK, OJ, OW, SB, LvW, TLi; Investigation: IT, TLe, HWK, MMH, SA, BM, SB, LvW, TLi; Visualization: IT, TLe; Writing – Original Draft: HMM, IT; Writing – review & editing: all the co-authors; Project administration: IT, HMM; Supervision: HMM, HWK, OJ, OW, JW, HS, JoW, JeW; Funding Acquisition, HMM.

## Acknowledgements

First, we thank Sonja Kachler, Gabriele Paetzold and Sandra Theil for the excellent technical assistance. Also, we would like to thank Prof. S. Weggen (Heinrich Heine University, Germany) to have kindly provided the IC16 Antibody. We thank Prof. H.-J. Knölker (Technische Universität Dresden, Germany) for providing the synthetic Aβ-3-40 peptide. We thank Niels Hansen and Anke Jahn Brodmann for support and for providing CSF-samples from the local biobank. H.M.M. acknowledge funding by the Interdisziplinäres Zentrum für Klinische Forschung (IZKF) of Würzburg, project number A-F-N-419. H.M.M. acknowledges the Junior Group Leader program of the Rudolf Virchow Center, the excellent ideas programme of the University of Würzburg and support through the Emmy Noether Programme of the Deutsche Forschungsgemeinschaft (DFG) (MA6957/1-1). T.L. acknowledges funding by the Graduate School of Life Sciences of the University of Würzburg. O.W. acknowledges funding by the DFG (WI3472/11-1 and Research Training Group 2824 (RTG2824)). J.W acknowledges funding by the DFG WA1477/6-6. M.M.H is a member of RTG 2824 funded by the DFG.

---

**Figure 1 - figure supplement 1.**
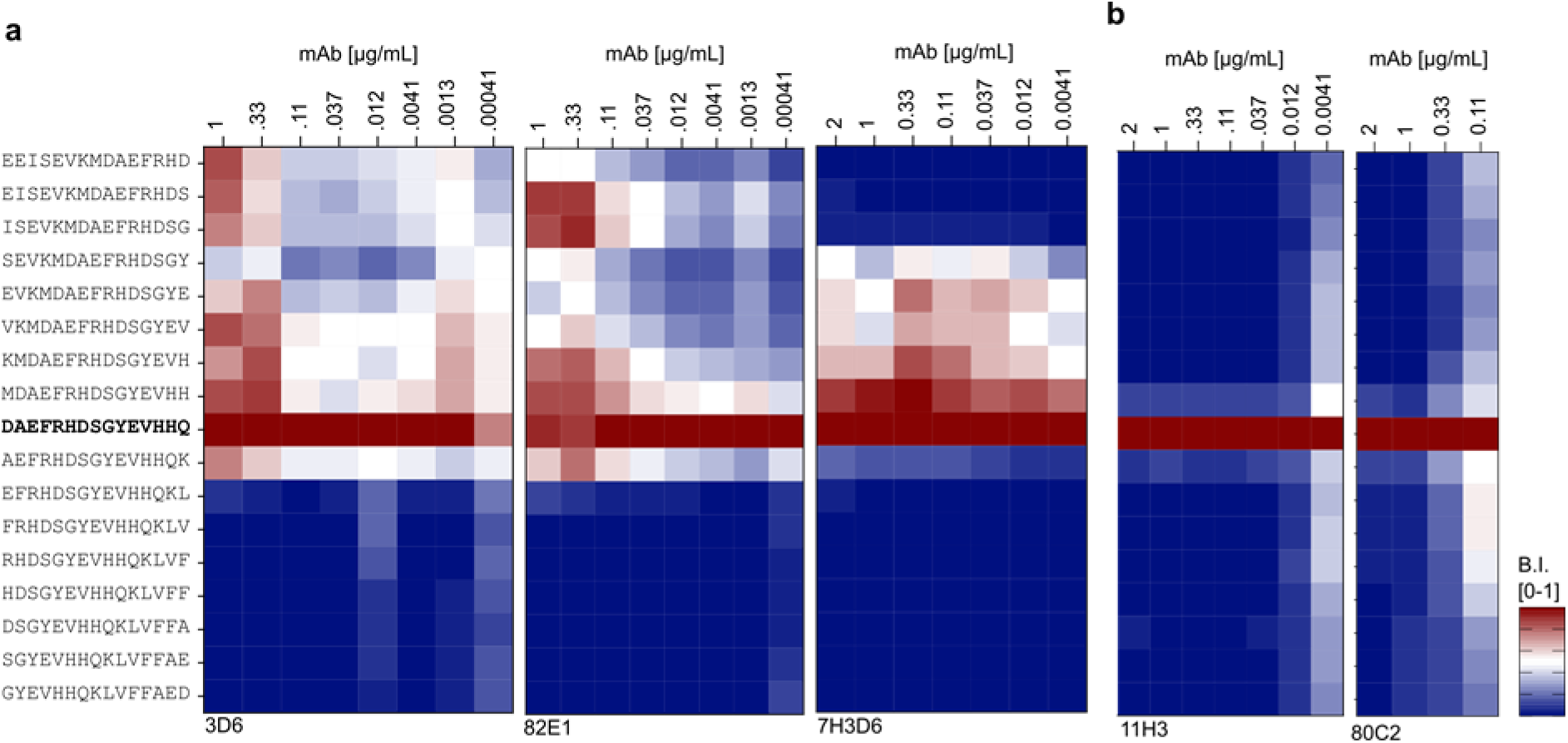
Titration series of N-terminal specific Anti-Aβs. (a) Titration of monoclonal Aβ-Ab 3D6, 82E1 and 7H3D6 at concentrations ranging from 2 to 0.00041 µg/mL to study the selectivity for free-Asp(1) over Asp(1) within an N-terminal elongated peptide. The signals were normalized internally and presented as a heat map with blue-to-red shades ranging from 0 to 1. (b) Monoclonal Aβ-Ab 11H3D6 and 80C2 exhibited high selectivity towards free-Asp(1) across the chosen concentration range of 2 µg/mL down to 0.0041 µg/mL.

**Figure 2 - figure supplement 1.**
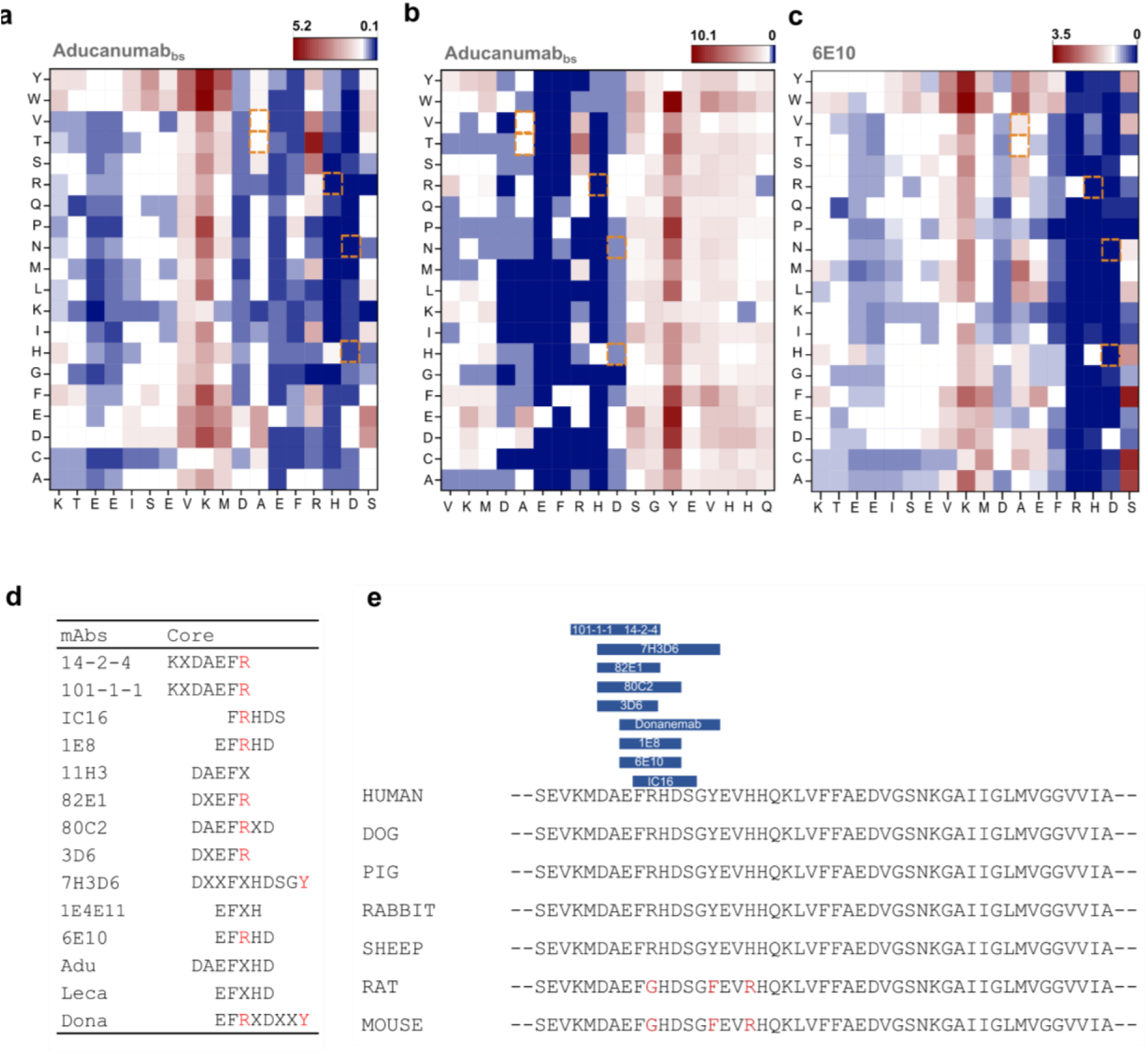
N-terminally elongated Deep Mutational Scans of Aducanumab_bs_ and 6E10 with core motifs summary. (a) Fingerprint analysis for the first peptide group covering the sequence *KTEEISEVKMDAEFRHDS*. The Aβ-Ab tested here is Aducanumab_bs._ (b) Fingerprint analysis for the second peptide group covering the sequence *VKMDAEFRHDSGYEVHHQ*. The Aβ-Ab tested here is Aducanumab_bs._ (c) Fingerprint analysis of 6E10 on the sequence *KTEEISEVKMDAEFRHDS*. (d) Summary of the antibody core motifs, “X” within the sequence stands for non-conserved residue. When an amino acid variation led to a decrease in binding below 50% of wt, it was marked as dominant negative. If more than 10 amino acid substitutions were dominant negative, the entire position was designated as part of the core motif (represented in single-letter coding). (e) Aβ-Ab species preferences. Amino acid positions which are different in human and rat/mouse are highlighted in red. These specific positions were probed. Antibodies showing altered binding strength after replacing R5 or Y10 in the human Aβ sequence by G5 or F10 (rat/mouse Aβ sequence) are highlighted in blue. Human: Homo Sapiens; Dog: Canis Lupus Familiaris; Pig: Sus Scrofa; Rabbit: Oryctolagus Cuniculus; Sheep: Ovis Aries; Rat: Rattus Norvegicus; Mouse: Mus Musculus.

**Figure 2 – figure supplement 2.**
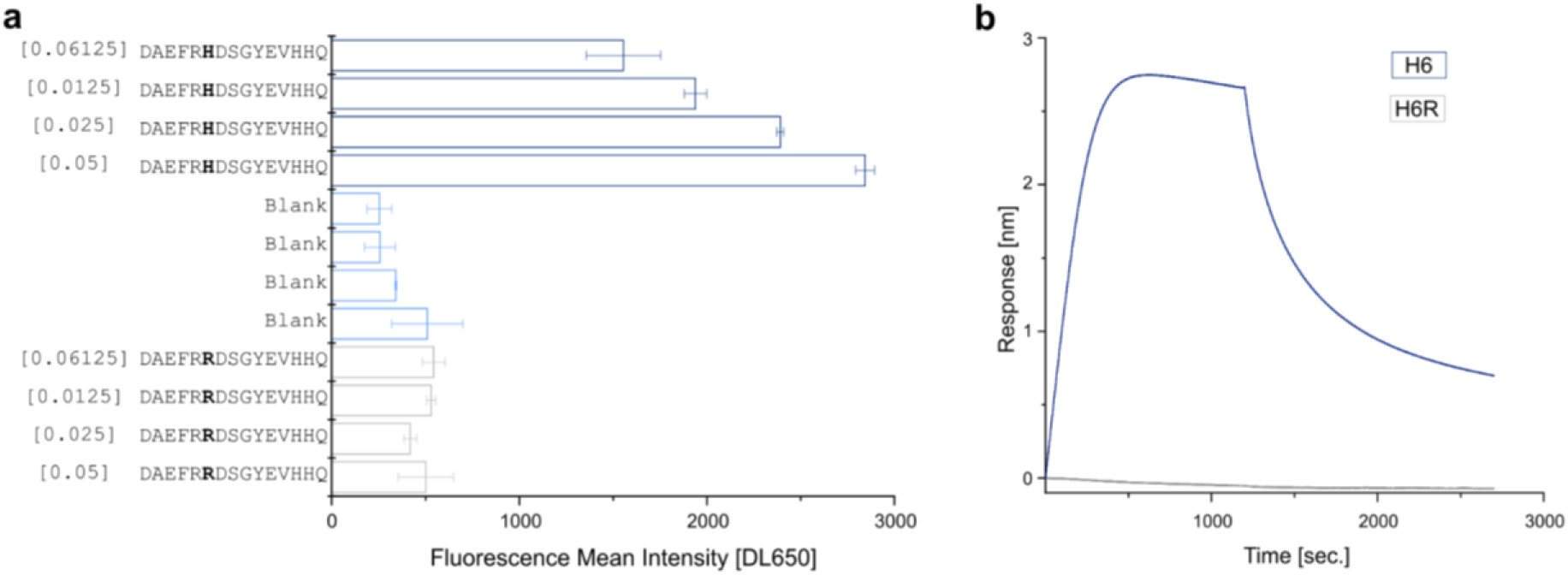
Aβ1-15 carrying the English mutation (H6R) shows reduced affinity for Aducanumab_bs._ (a) Dot Blot from the purified 6xHis peptides *DAEFR**H**DSGYEVHHQ* and *DAEFR**R**DSGYEVHHQ* were printed using Slide Spotter robot. IgG signals were detected using Goat anti-human DL650, in square brackets the µg/mL of Aducanumab_bs._ (b) The Biolayer interferometry responses of Aducanumab_bs_ (10nM) over the NTA-loaded *DAEFR**H**DSGYEVHHQ* and *DAEFR**R**DSGYEVHHQ* peptides are shown.

**Figure 3 – figure supplement 1.**
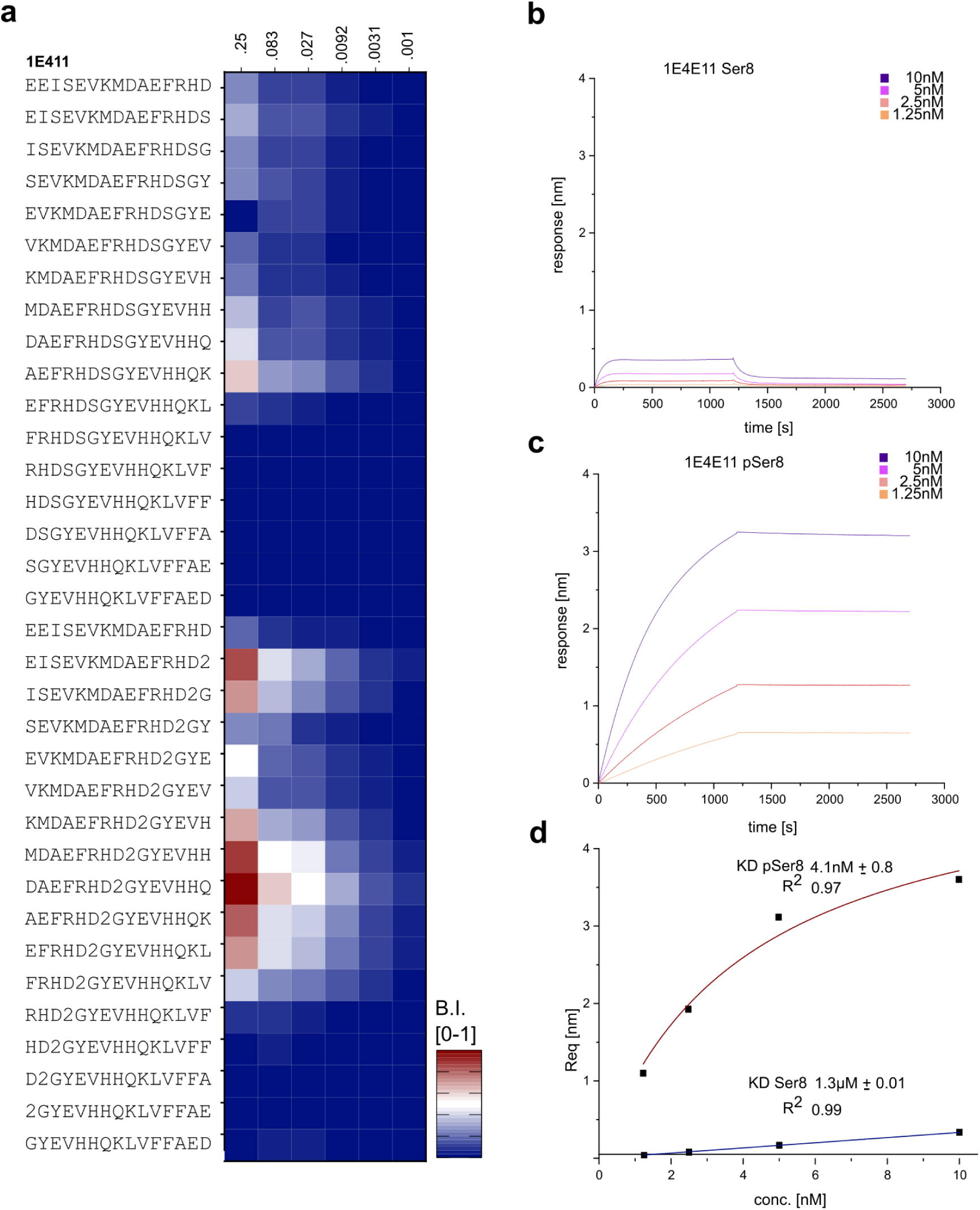
Antibody 1E4E11 shows high preference for phosphorylated pSer8 Aβ1-15 in µSPOT and BLI assays. (a) Titration of monoclonal Aβ-Ab 1E4E11 at concentrations ranging from 0.25 to 0.001 µg/mL to achieve high selectivity for Aβ1-15 phosphorylated at Ser(8). The modified pSer is represented as ’2’. The signals were globally normalized against *DAEFRHDpSGYEVHHQ* and presented as a heat map with blue-to-red shades ranging from 0 to 1. (b) BLI dose-response curve of mAb 1E4E11 binding to the NTA-loaded *DAEFRHDSGYEVHHQ-6xHis-amide* peptide. (c) Biolayer interferometry dose-response curve of mAb 1E4E11 binding to the NTA-loaded *DAEFRHDpSGYEVHHQ-6xHis-amide* peptide. (d) BLI steady-state analysis revealed K_D_s of 4.1nM and 1.3µM for Aβ1-15 with phosphorylated and unphosphorylated Ser (8), respectively (1:1 model).

**Figure 3 – figure supplement 2.**
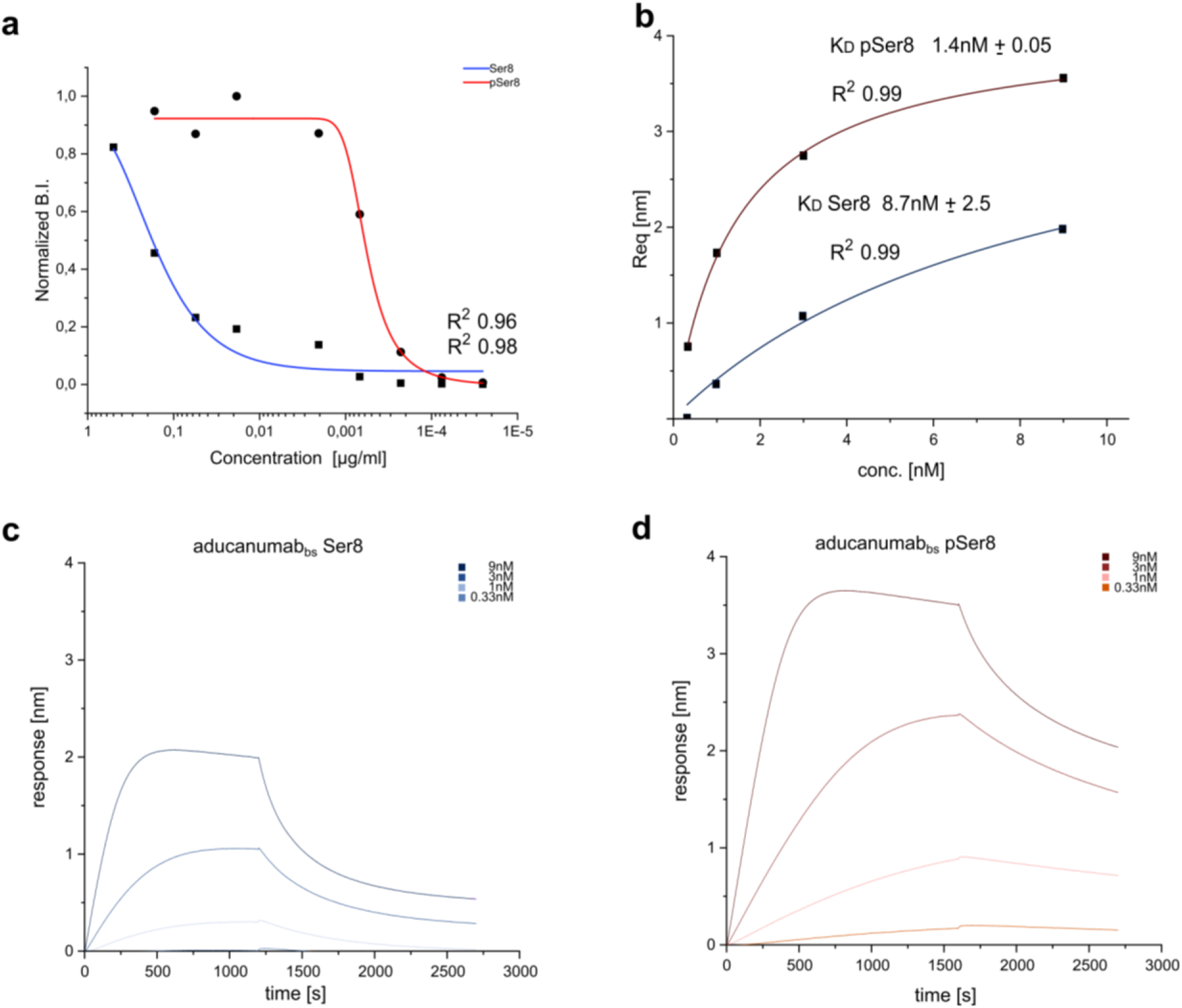
Phosphorylation aids Aducanumab_bs_ binding to synthetic monomeric Aβ1-15. (a) Titration of Aducanumab_bs_ at different concentrations to achieve high selectivity for phosphorylated pSer8-Aβ1-15 over unphosphorylated Aβ1-15. The signals were internally normalized and fitted as a sigmoidal curve (Boltzmann). The phospho signal for pSer8-Aβ1-15 is shown in red (r^2 0.98) and the signal for unphosphorylated Aβ1-15 is shown in blue (r^2 0.96). (b) BLI steady-state analysis revealed KDs of 1.4nM and 8.7nM for phosphorylated and unphosphorylated Aβ1-15, respectively (1:1 model). (c) Biolayer interferometry dose-response curve of Aducanumab binding to the NTA-loaded *DAEFRHDSGYEVHHQ-6xHis-amide* peptide. (d) Biolayer interferometry dose-response curve of Aducanumab binding to the NTA-loaded *DAEFRHDpSGYEVHHQ-6xHis-amide* peptide.

**Figure 3 – figure supplement 3.**
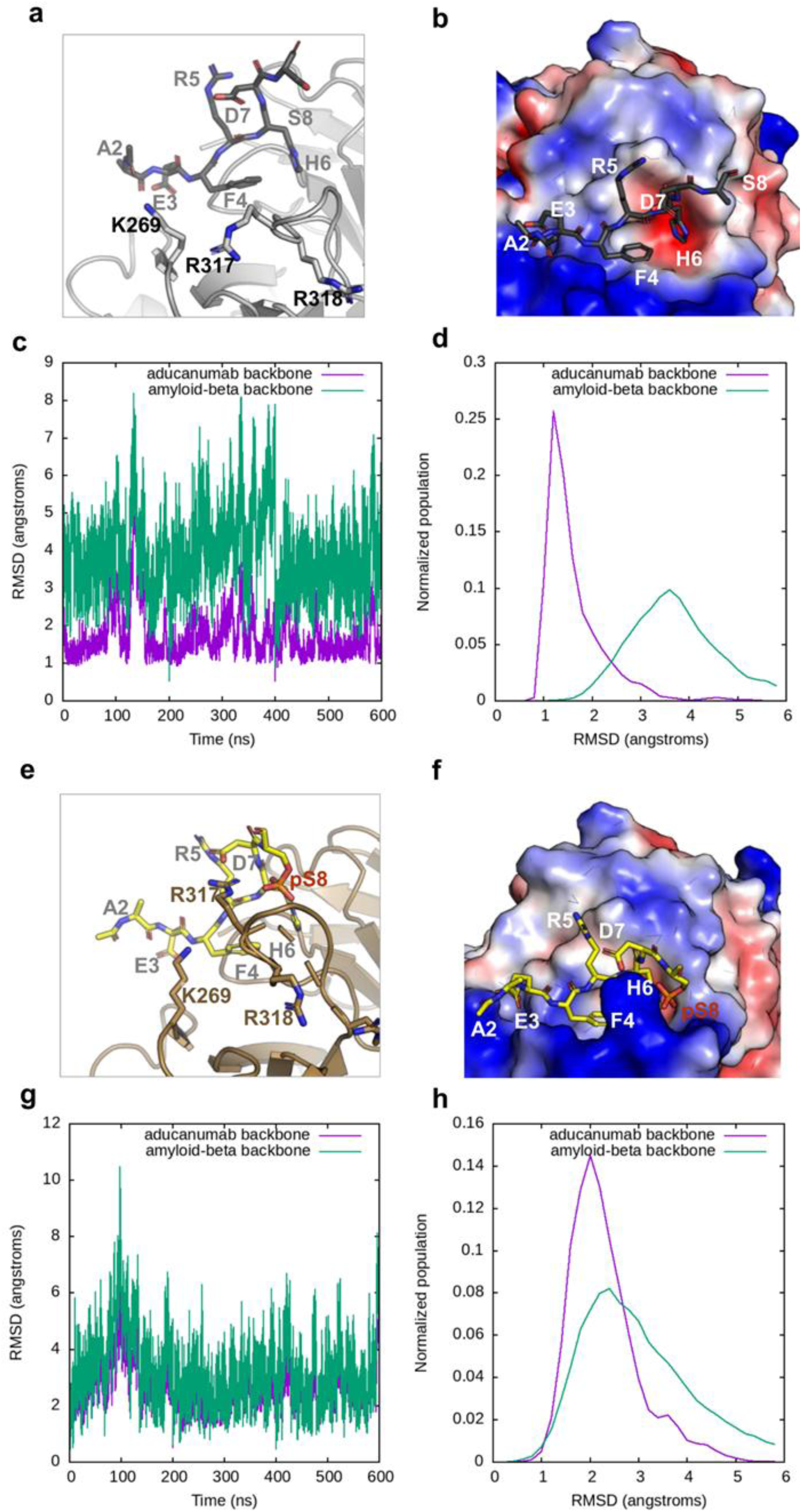
MD simulation of Aducanumab with pSer(8)-Aβ2-8. (a) Sticks and cartoon depiction of the MD simulations between Aducanumab and AEFRHDS. (b) Electrostatic density for the non-phosphorylated sequence interaction. (c-d) RMSD analysis of the backbone atoms of the unphosphorylated candidate (average conformation over 600ns). (e) Sticks and cartoon depiction of the MD simulations between Aducanumab and AEFRHDpS. (f) Electrostatic density for the pSer8 sequence interaction. (g-h) RMSD analysis of the backbone atoms of the PTM candidate (average conformation over 600ns).

**Figure 3 – figure supplement 4.**
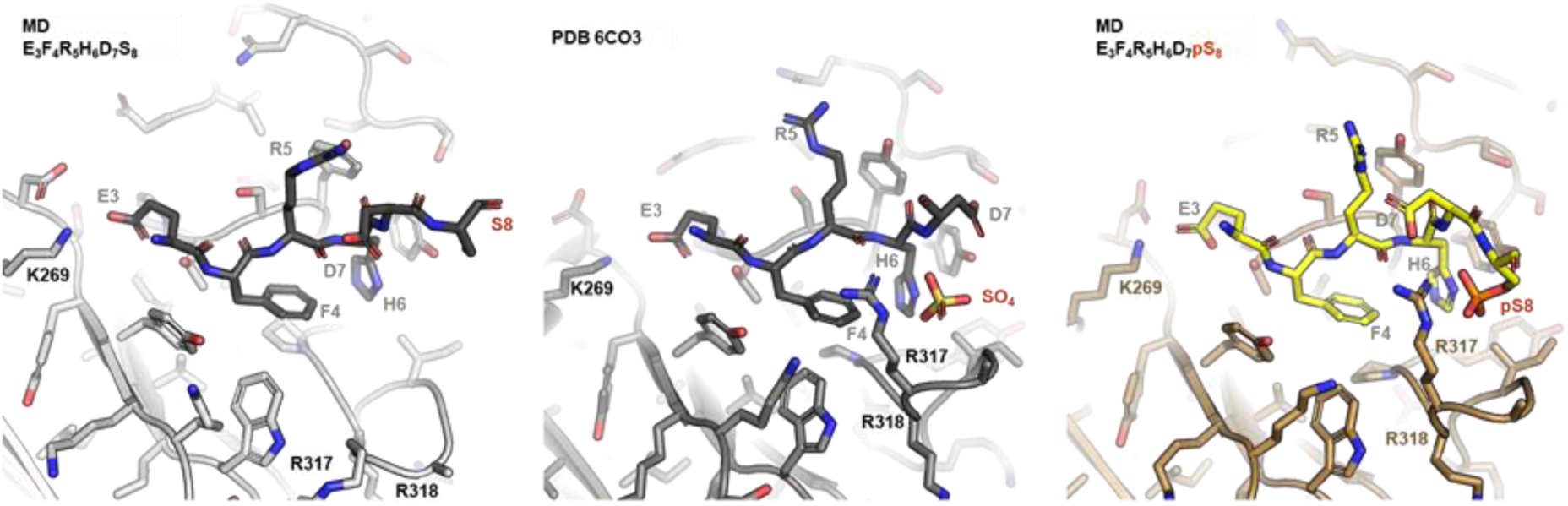
pSer8 but not Ser8-Aβ recapitulate the binding mode resolved by christallography. Comparison of MD simulations with structural data. The MD simulation was conducted with the resolved epitope peptide *EFRHDS,* but in the absence of additional ions, and predicts a largely altered binding mode characterized by over 10 Å movements of R317 and R318. In contrast, the crystal structure (PDB 6CO3) harbors a sulfate ion that neutralizes the positive charge introduced by R317 and R318. Strikingly the MD of the pSer(8) modified peptide recapitulates the resolved X-ray binding mode to monomeric Aβ. Aducanumab targets aggregated Aβ, while still engaging with low micromolar affinity a monomeric Aβ.

**Figure 6 – figure supplement 1.**
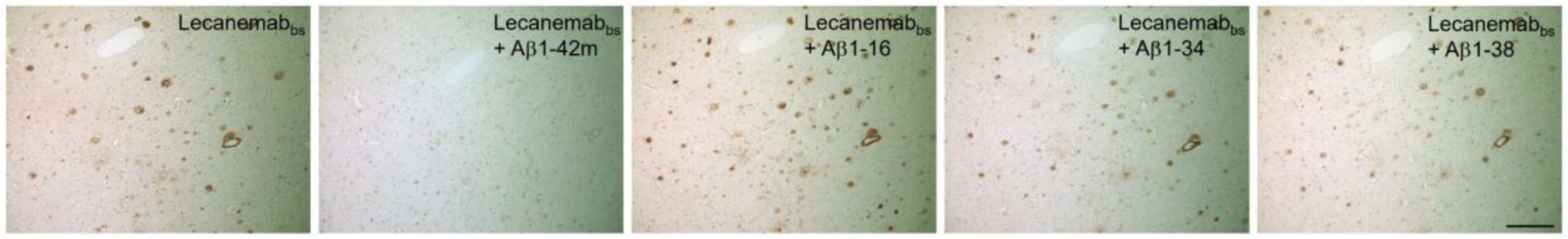
HFIP treated Aβ1-42, but not shorter synthetic Aβ-peptides block Lecanemab_bs_ binding to amyloid plaques. Parallel sections from an AD patient were stained, from left to write, with Lecanemab_bs_ only (BioXCell #BXC-SIM0032) or pre-adsorbed with either Aβ1-42 monomers (Aβ1-42m), Aβ1-16 (Aβ1-16), Aβ1-34 (Aβ1-34) and Aβ1-38 (Aβ1-38). Scale bar: 200 µm.

